# Advanced in silico validation framework for three-dimensional Traction Force Microscopy and application to an in vitro model of sprouting angiogenesis

**DOI:** 10.1101/2020.12.08.411603

**Authors:** J. Barrasa-Fano, A. Shapeti, J. de Jong, A. Ranga, J.A. Sanz-Herrera, H. Van Oosterwyck

## Abstract

In the last decade, cellular forces in three-dimensional hydrogels that mimic the extracellular matrix have been calculated by means of Traction Force Microscopy (TFM). However, characterizing the accuracy limits of a traction recovery method is critical to avoid obscuring physiological information due to traction recovery errors. So far, 3D TFM algorithms have only been validated using simplified cell geometries, bypassing image processing steps or arbitrarily simulating focal adhesions. Moreover, it is still uncertain which of the two common traction recovery methods, i.e., forward and inverse, is more robust against the inherent challenges of 3D TFM. In this work, we established an advanced in silico validation framework that is applicable to any 3D TFM experimental setup and that can be used to correctly couple the experimental and computational aspects of 3D TFM. Advancements relate to the simultaneous incorporation of complex cell geometries, simulation of microscopy images of varying bead densities and different focal adhesion sizes and distributions. By measuring the traction recovery error with respect to ground truth solutions, we found that while highest traction recovery errors occur for cases with sparse and small focal adhesions, our implementation of the inverse method improves two-fold the accuracy with respect to the forward method (average error of 23% vs. 50%). This advantage was further supported by recovering cellular tractions around angiogenic sprouts in an in vitro model of angiogenesis. The inverse method recovered more realistic traction patterns than the forward method, showing higher traction peaks and a clearer pulling pattern at the sprout protrusion tips.

## 1. Introduction

Hydrogel substrates are typically used to mimic the extracellular matrix (ECM). Over the last decades, manipulation of ECM properties has revealed that cell fate decisions, (such as migration, proliferation, differentiation or death) are modulated by a cell’s mechanical microenvironment, which is crucial for single cell behavior as well as multicellular organization ([1, 2, 3, 4, 5, 6]). Stem cell fate can be directed by ECM stiffness ([3, 7, 8, 9, 10, 11]), stress relaxation properties ([12]) or its adhesive ligands ([13]). ECM stiffness can also regulate malignancy of tumor cells ([4, 14, 15, 16, 17, 18]), cell contraction ([19]), and cell spreading ([20, 21]). Cells probe and adapt their mechanical environment by actively exerting forces on neighboring cells and on the ECM. As these forces are essential for the understanding of mechanotransduction, there is a strong need for methods that measure them accurately. Cellular mechanical interactions with the ECM are typically measured by means of traction force microscopy (TFM) ([22, 23]). Briefly, fiducial markers (such as fluorescent beads) are normally embedded into hydrogels of known mechanical properties. Microscopy imaging is used to acquire a hydrogel deformed or *stressed* state (with mechanically active cells) and a hydrogel stress-free or *relaxed* state (obtained before cell seeding or after disruption of a cell’s ability to apply force). Then, hydrogel deformations caused by the cellular forces are measured using image processing. Stress, tractions or forces can be inferred from the deformations, given the mechanical properties of the hydrogel. While TFM has been widely used for the last 20 years in 2D cell cultures to compute the exerted traction fields on planar substrates, measuring cellular forces in 3D is increasingly viewed as more physiologically relevant ([24]). However, extending TFM to 3D entails the following problems:

Cells use integrins to bind to ECM proteins at discrete areas called focal adhesions (FAs) ([25]). Through FAs, the connection between ECM and the mechanically active acto-myosin cytoskeletal network, is formed ([26]). While FAs in 2D are typically punctuated and discrete, there is still uncertainty of their size and distribution in 3D. On the one hand, some studies reported diffusely distributed FAs over the cell membrane [27, 28]. On the other hand, others have shown more discrete patterns varying from less than 1*μ*m up to 4*μ*m [29, 30, 31, 32, 33]). The magnitude and distribution of the tractions depend on the size of the areas over which cellular forces are applied. Similarly, the frequency content of the displacement field in the ECM is also given by the size and the density of the FAs. This is a major concern for the selection of the bead density, which determines the spatial sampling frequency. According to the Nyquist sampling theorem, spatial sampling frequencies should be higher than twice the maximum frequency of the displacement field in the ECM ([34]). However, the use of high bead densities in 3D TFM is not possible in practice. Too high bead densities can hamper their tracking due to overlapping caused by the point spread function of the microscope and the mechanical properties of the hydrogel or cell behavior might change. Therefore it is clear that it critical to determine and optimize the relation between 3D TFM accuracy with respect to bead density, FA size and density is vital for a correct coupling of the experimental and the computational 3D TFM workflows. However, this problem has been largely overlooked in the field of TFM. The measured displacements, regardless of the algorithm used (e.g. Particle Tracking ([35, 36]), Particle Image Velocimetry (PIV) ([37, 38]) or Free Form Deformation-Based Image Registration (FFD) ([39]), thus inherently contain sampling noise that hampers traction recovery. Moreover, it is still uncertain which of the two main traction recovery methods in literature (i.e. the forward method and the inverse method) is more robust against these inherent limitations:

### The forward method

The terminology used for this method can often be misleading in the literature. For consistency, we follow the terminology used in [40, 41]. This method can be found in two different forms in the literature depending on how the stress tensor is stated: (i) in its strong form and hence using numerical derivatives, (ii) in its weak form and hence using finite element (FE) analysis. Numerical derivatives are used to calculate the strain tensor from the measured displacement field and the stress tensor through the constitutive law of the hydrogel [40, 41, 42]. Alternatively, the stress tensor can be calculated by means of a FE solver. A FE mesh of the cell surface and the hydrogel is created and the measured displacements are defined as an input condition at the nodes of the mesh ([43, 44, 45]). Regardless of the way the stress tensor is calculated, cellular tractions are obtained by means of Cauchy’s stress formula and the cell surface normal vectors. However, the sampling noise present in the *measured* displacements might propagate through traction recovery ([46]).

### The inverse method

This approach minimizes the difference between the measured displacements (obtained analogously to the forward method) and a mathematically consistent (regularized) solution. While the most common way of solving this problem is by minimizing a least square estimate with Tikhonov regularization ([35]), alternative inverse formulations have recently been developed ([47, 48, 49, 50]). We recently proposed an inverse method that fulfills the equilibrium of internal forces with real acting forces, which is not always warranted in other inverse methods ([51]).

Existing works, typically validate *in silico* 3D traction recovery algorithms by simulating ground truth displacements and tractions and by comparing them with the results obtained by the algorithm. However, they often simplify or bypass critical aspects, i.e., cellular geometries, FAs and displacement sampling noise. Authors have used simplified geometries (instead of real cell geometries) that ease calculations such as cylindrical ([48]), spherical ([52]) or ellipsoidal inclusions ([53]). Recently, real cell geometries obtained from 3D optical microscopy imaging (such as confocal imaging) have been used ([54, 49, 50]). However, these studies use a single arbitrarily chosen FA distribution and size. So far, no studies have reported validations systematically varying FA size or distribution. Furthermore, displacement sampling noise is often simulated by adding stochastic noise to the ground truth displacement field ([34, 48, 51]). Other studies use a more realistic approach by simulating microscopy bead images as it incorporates the inherent sampling noise [39, 55, 52]. Again, these studies either do not use real cell geometries, or do not systematically vary FA size and distribution.

With this study, we aim to underline the importance of selecting a traction recovery method that can overcome the inherent limitations of 3D TFM and characterizing its accuracy limits. First, we refined the 3D TFM in silico validation framework by incorporating different FA sizes and distributions to generate ground truth displacements and tractions and by simulating image stacks accounting for diffraction limited optical microscopy. We thereby establish a more realistic and rigorous validation framework that allows us to investigate the relation between bead density and FA size and distribution with 3D TFM accuracy, which is critical for a correct coupling of the experimental and computational steps of TFM. Second, we compared the accuracy of the forward and our inverse traction recovery method, with respect to the ground truth data, and proved the superiority of the latter. We then performed an experimental study for sprouting angiogenesis in a polyethylene glycol (PEG) hydrogel, and applied both traction recovery methods. Again, our inverse method showed more realistic traction patterns further supporting our in silico results.

We chose sprouting angiogenesis as an application for both our in silico simulations and for our in vitro experiments. On the one hand, we have previously established experimental and computational protocols to measure 3D hydrogel displacements during endothelial cell invasion in an in vitro model of sprouting angiogenesis [56]. On the other hand, angiogenic sprouts typically show multiple branches and cellular protrusions [56]. The increased complexity of these geometries compared to those of single cells, represents a more interesting and challenging case for assessing accuracy in 3D TFM.

## 2. Methods

We will first describe the different steps that are taken to evaluate the accuracy of 3D traction recovery algorithms (see a general overview in Fig. 1). The evaluation is performed for a real 3D sprout geometry that was acquired by means of confocal microscopy (see Appendix A for a brief summary of the experimental model). Ground truth traction and displacement data was generated by creating a 3D confocal image-based finite element model (section 2.1) for the selected sprout and for cellular tractions that lead to displacement fields qualitatively similar to the ones we measured before ([56]). The simulated displacement fields serve as input for the simulation of microscopy images of bead distributions (before and after the application of cellular tractions; section 2.2) that are used as input for the 3D TFM workflow (and that involve displacement measurement and traction recovery, section 2.3). The calculated displacements and tractions are then compared to the ground truth displacement and tractions (for both the forward and inverse method) and proper error metrics (section 2.4) are defined to evaluate the accuracy of both traction recovery methods.

**Figure 1:**
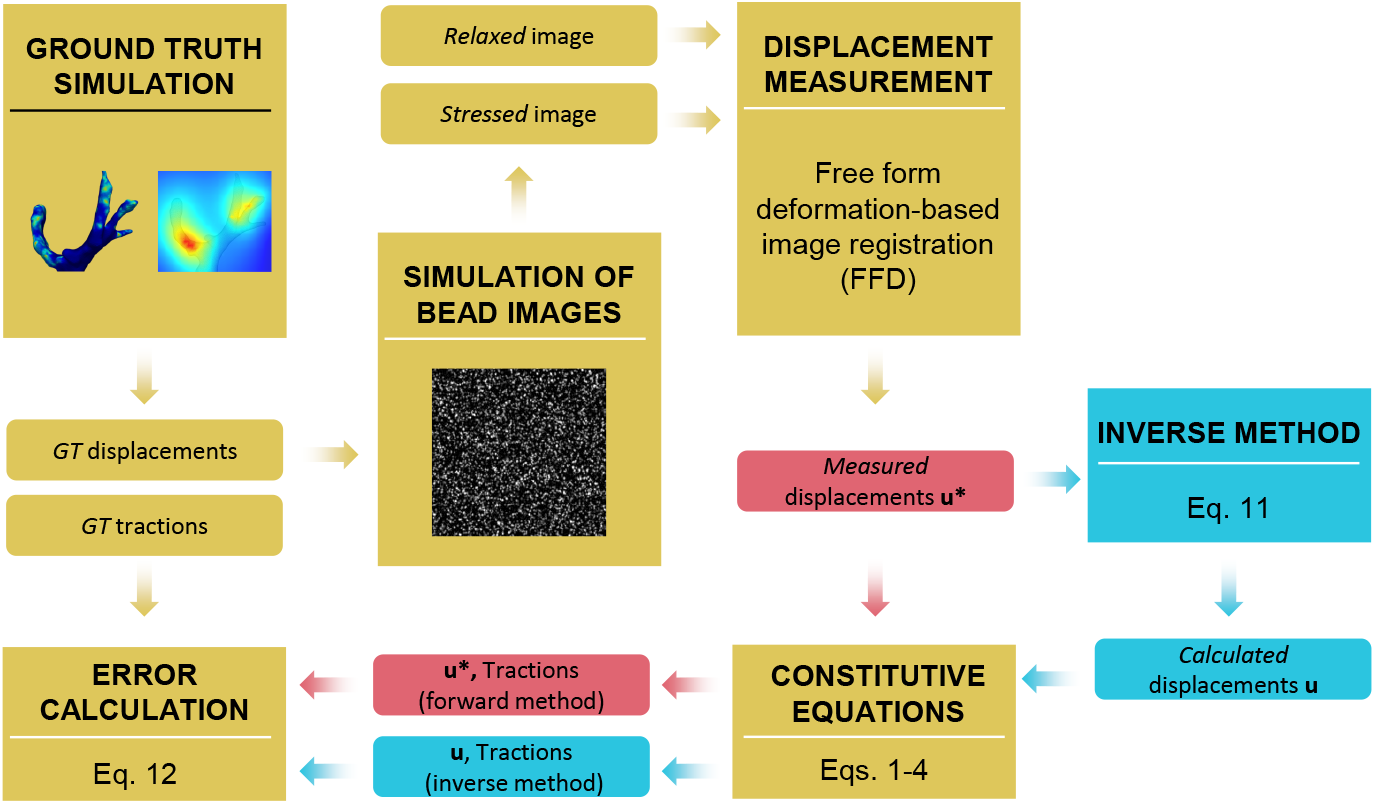
Workflow used for ground truth (*GT*) simulations and for validation of the forward (red) and inverse (blue) traction recovery methods. Common steps for both methods are depicted in yellow. The figure refers to the key equations in the text.

We then apply both traction recovery methods to a new data set of sprouting angiogenesis in a PEG hydrogel that was acquired as part of this study (making use of the same in vitro setup as described in Appendix A) and that will illustrate the importance of selecting a proper method for the accuracy of traction recovery. A PEG hydrogel was used here because of its linear elastic behavior [35] and because we evaluated TFM accuracy for ground truth simulations that assumed linear elastic hydrogels (although both the forward and inverse methods are compatible with nonlinear elasticity, see also [51]).

### 2.1. Ground truth simulations

The image stack of an angiogenic sprout (see Fig. 2a) was denoised using penalized least squares-based background ([57]) and enhanced by a contrast stretching operation. Subsequently, a cellular surface segmentation was thresholded applying Otsu’s binarization algorithm ([58]). We then segmented the four main protrusions of the sprout (see Fig. 2b) to define the areas where focal adhesions are simulated. The principal direction of each protrusion was computed to later define the direction of the ground truth tractions (see details in Appendix B). We then used the Matlab toolbox Iso2Mesh [59] to create a surface mesh of the sprout and a volumetric tetrahedral mesh of the hydrogel volume (see Fig. 2c). The inner surface of the hydrogel volume was defined by the sprout surface and the outer surface was a 107*×*111*×*40 *μm* cube. The mesh had around 173000 4-node tetrahedral elements.

**Figure 2:**
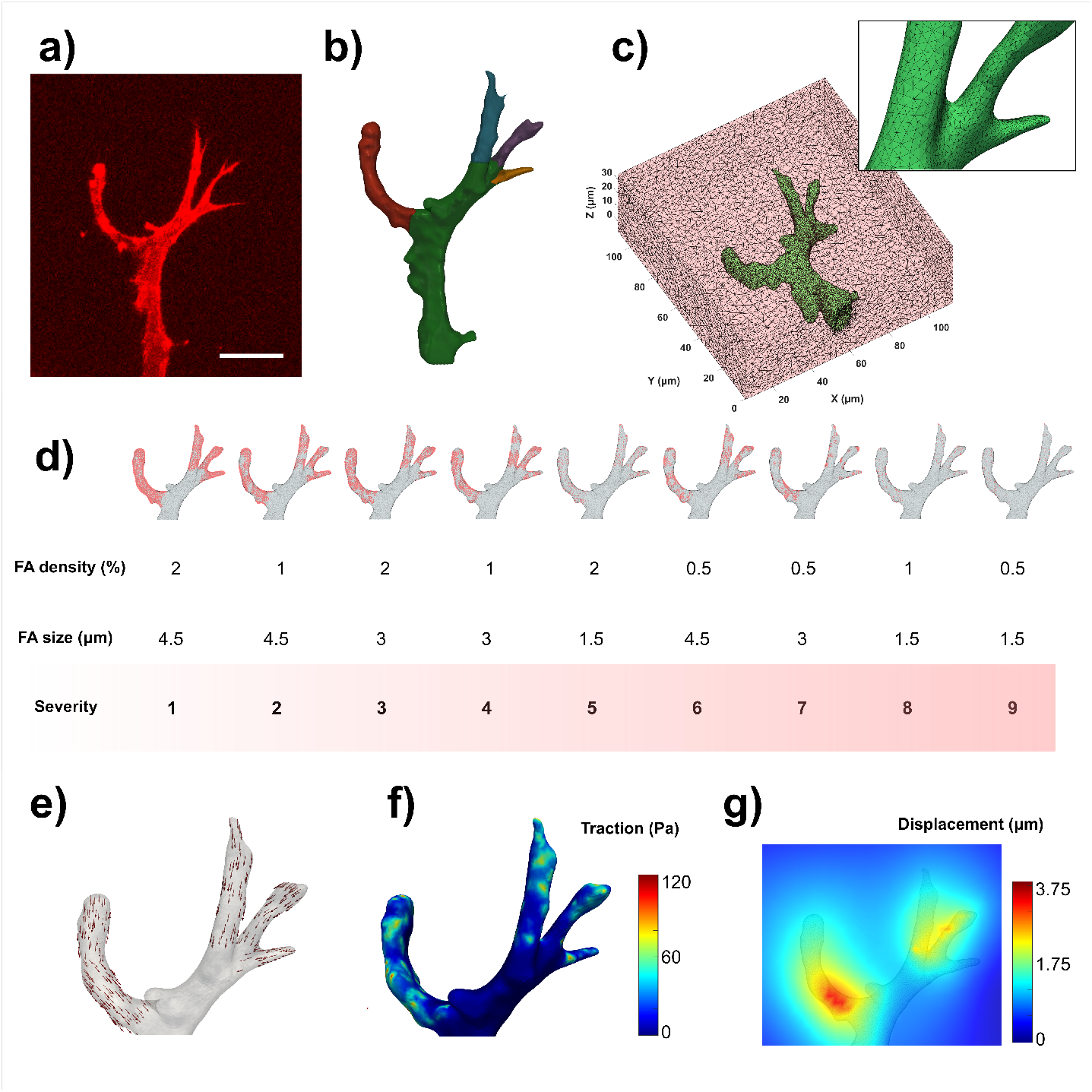
Ground truth generation. (a) Maximum intensity projection of the image stack acquired by means of confocal microscopy of the angiogenic sprout used for the simulations. Scale bar: 21 *μ*m. (b) Sprout segmented geometry in green and the four segmented protrusions in red, blue purple and orange. (c) Finite element mesh including the sprout geometry (green) and the hydrogel domain (pink) and a close up of the sprout surface faces. (d) Systematic variation of focal adhesion size and distribution. Focal adhesion nodes (red) are depicted over the rest of the finite element mesh (gray). (e-g) Example of ground truth generation for severity case 5. Nodal forces (e) are applied on the focal adhesion nodes (magnitude: 246.48 pN) and ground truth tractions (f) and displacements (g) are obtained.

Given the current uncertainty on the size and distribution of FAs in 3D, we systematically modeled different kinds of FA patterns, from relatively discrete, to relatively continuous (see Fig. 2d). FAs were defined on the nodes of the surface of the four sprout protrusions according to:

- Focal adhesion density: percentage of protrusion nodes randomly selected to conform the central node of a FA. In this study we considered: 0.5%, 1% and 2%.
- Focal adhesion size: mean distance between the central FA node and the nodes on the outer perimeter of the FA. In this study we considered: 1.5, 3 and 4.5 *μ*m.

Nodal forces in the direction of the sprout branches were then prescribed on FA nodes to mimic typical pulling patterns shown in literature [48, 56] (see Fig.2e). The sum of all the nodal forces was constant for all the cases and fixed at 70nN (see details in Appendix C). This value was calibrated to obtain a corresponding displacement field of an order of magnitude similar to previous works ([35]). Then, ground truth (*GT*) tractions and displacements expressed in the FE mesh nodes were obtained (see Fig. 2f,g). Nine different *GT* cases were run that corresponded to 9 different combinations of FA density (3 values) and size (3 values). We defined a severity index (from 1 to 9) that describes the heterogeneity (non-smoothness) of the traction field and that was associated to the coefficient of variation (COV) according to the formula: 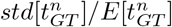, where 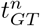 is the value of the magnitude of the ground truth traction at the *n* nodes of the sprout boundary domain and std[ ] and E[ ] stand for standard deviation and mean, respectively (see Fig. C.9). The 9 cases were ranked according to increasing values of traction field COV and a severity index was assigned to each of them, with 1 for the lowest COV (and therefore smoothest traction field) and 9 for the highest COV (least smooth traction field). A case (and associated traction field) with a high severity index will give rise to a displacement field with a high frequency content and will therefore lead to more sampling errors for a given bead density. The hydrogel domain was modeled as a linear elastic material with an elastic modulus of 200 Pa and a Poisson’s ratio of 0.3 [–]. These cases were run using Matlab R2019a and Abaqus Simulia 6.14 (see [51] for further details of the implementation).

### 2.2. Simulation of microscopy bead images

To assess the accuracy of traction recovery methods under conditions that are representative for a real TFM experiment, we applied the ground truth displacements to synthetically generated microscopy bead images. Diffraction patterns were simulated using a point-spread function (PSF) determined by fitting a 3D Gaussian profile to 4000 beads from a real experimental image (more details are provided in Appendix D). The intensities of these PSFs were imposed at random locations of the image (see Figure 3a). Five different experimentally feasible bead densities were considered, namely: 0.005, 0.01, 0.03, 0.05 and 0.07 beads/*μm*^3^ (see Figure 3b and Table D.2). Moreover, since the simulated beads were randomly distributed, 20 realizations were considered for each bead density to provide reliable performance results. Similarly, *stressed* state images were generated, by shifting each bead location according to the ground truth displacement field (interpolated from the FE nodes to the image coordinates).

**Figure 3:**
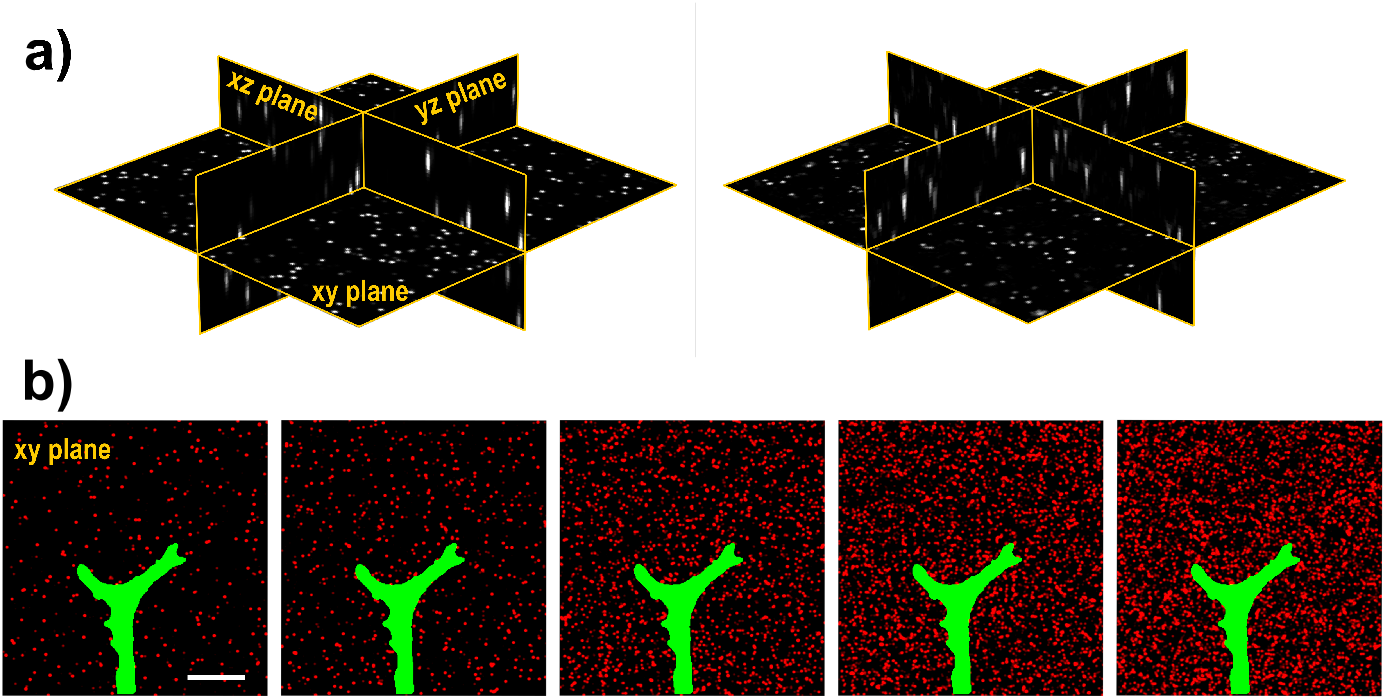
Simulation of microscopy bead images. (a) Central slice planes of a simulated (left) and a real (right) 0.2 micron-sized fluorescent spheres image stack acquired by means of confocal image microscopy. Volume size is 107*×*111*×*40 *μm*^3^. (b) Central xy plane (sectioning plane of the confocal microscope) of simulated bead images with bead density from 0.005 to 0.07 (left to right) beads/*μm*^3^. Beads are depicted in red and the sprout geometry in green for visual reference. Scale bar: 22 *μ*m.

### 2.3. Traction field recovery

First, B-spline-based FFD ([39]), a method that showed higher accuracy compared to PIV, was used to measure displacement fields from the *relaxed* and the *stressed* images (more details on the parameters used can be found in Appendix E). The displacement field values were interpolated from the image coordinates to the FE nodes. From here onward, they are referred to as *measured* displacement field, **u**^*^ (see Fig. 1). Second, tractions were recovered using the forward and inverse methodologies in the framework of the Finite Element Method (FEM). The analysis is performed assuming a linear elastic behavior of the hydrogel as detailed in the next section.

#### Forward method

From the *measured* displacement field **u**^⋆^, tractions are straightforwardly obtained following the FEM procedure (see the red arrows in Fig. 1). First, the strain field is computed as,

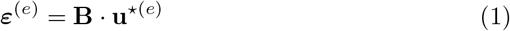

where *ε*^(*e*)^ is the strain tensor field computed at each Gauss point of element (*e*), **B** the gradient matrix of the shape functions, and **u**^*(*e*)^ the vector components of the *measured* displacement field at the nodes of the referred element. On the other hand, stresses are computed by means of the (linear, elastic) constitutive behavior,

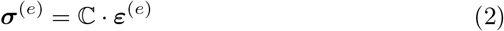

ℂ being a fourth order tensor which contains the elastic properties of the hydrogel, and ***σ***^(*e*)^ is the stress tensor field computed at each Gauss point of element (*e*). If an isotropic and homogeneous behavior is assumed, Eq. (2) yields,

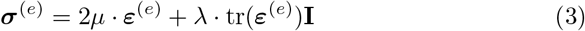

where *μ* and *λ* are the Lamé properties of the material. Finally, tractions are computed at the boundary of the sprout domain using Cauchy’s stress theorem:

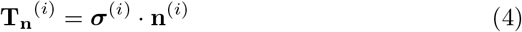

with **T**_**n**_^(*i*)^ the traction vector components at node *i* belonging to the sprout boundary, ***σ***^(*i*)^ the stress tensor field averaged at node *i* and **n**^(*i*)^ the outward normal to node *i*.

#### Inverse method

There are two main ingredients in the method: (i) Search for an inverse displacement field solution **u** as close as possible to the *measured* one **u**^⋆^, and (ii) which fulfills equilibrium of forces in the hydrogel domain ([51]). In a FEM framework, the discretized elasticity problem turns into the following algebraic system to solve,

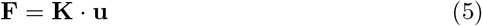

**F** is the nodal reaction forces vector which accounts for external or internal prescribed forces or displacements constraints at the nodes. If sprout boundary domain (s) and hydrogel domain (h) nodes are distinguished, Eq. (5) is reordered in the following way:

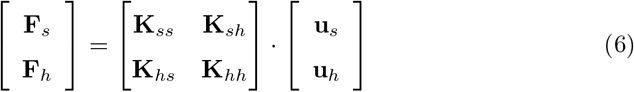

The ideas behind the inverse method exposed above, can be mathematically described as follows,

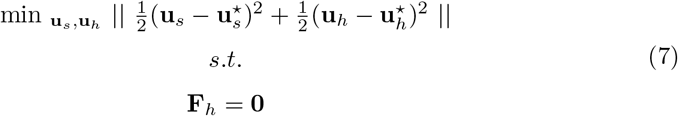

Note that this method is analogous to the so-called adjoint method in the inverse problem community. Further derivations of the inverse method to the analysis of TFM for linear elastic 2D materials can be found in [60, 61]. Using (6) in (7) yields,

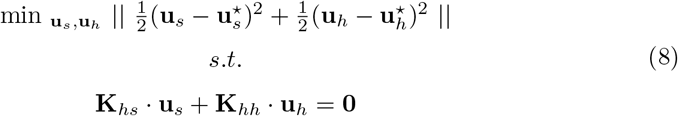

The equilibrium constraint in (8) can be included as a Lagrange’s multiplier (penalty) as follows,

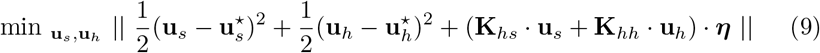

where ***η*** is the nodal-valued Lagrange multiplier scalar in the FE discretization.

The minmum of Eq. (9) can be determined analytically by means of Eq. (10):

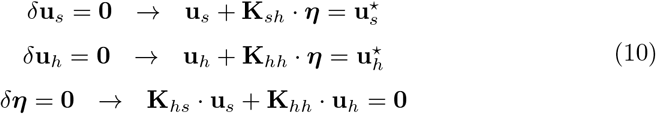

Eq. (10) can be written in matrix form as follows,

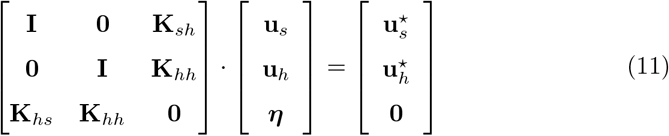

The solution of (11) provides the *calculated* displacement field **u** by means of th inverse method. The rest of the variables of the mechanical problem (strain, stress and traction) can be obtained using eqs. (1)–(4) substituting the *measured* forward displacement field **u**^⋆^ by the *calculated* inverse displacement field **u**(see the blue arrows in Fig. 1).

### 2.4. Error calculation

To compare the accuracy of the forward and the inverse method we define the following error metric:

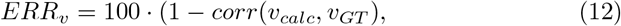

where *corr* is the linear correlation coefficient, *v*_*calc*_ and *v*_*GT*_ are the magnitude of the *calculated* and *GT* displacements or tractions at the nodes of the sprout boundary, respectively. Note that for the forward method, the *calculated* displacements are equal to the *measured* ones. Errors were computed for each of the analyzed cases (20 realizations *×* 5 bead densities *×* 9 severity cases) and plotted in boxplots in section 3.1. Additionally, the angle deviation of traction vectors was also calculated (see Appendix G).

## 3. Results

In this section, the errors obtained for both traction recovery methods with respect to the ground truth in silico simulations are shown. Additionally, we assessed the performance of both methods performed 3D TFM on real in vitro experimental data.

### 3.1. In silico simulations

Our *in silico* simulations show the strong effect of FA size and distribution and their associated traction field heterogeneity (as captured by the severity index) on 3D TFM accuracy. The increase in severity (from 1 to 9) leads to 5-10 times higher errors in displacement calculation (Fig. 4a: average errors range between 1 and 10% for both methods for a given bead density) and 2-3 times higher errors in traction calculation (Fig. 4b: average errors range between 40 and 70% for the forward method and between 15 and 55% for the inverse method). Increasing bead density (from 0.005 to 0.07 beads/*μm*^3^) overall reduces the error up to 10%. While both the forward and the inverse methods achieve relatively low errors in displacement calculation (below 10% for all the cases; Fig. 4c), a substantial difference can be seen in traction calculation. Traction errors are twice as high for the forward method as for the inverse method (median values of 50% and 23%, respectively; Fig. 4d). While the inverse method outperforms the forward method for all severity cases, the traction errors for the highest severity cases (7 to 9) are non-negligible for both methods, with average errors ranging from 50-62% (severity index 7; values correspond to highest and lowest bead density respectively) to 66-70% (severity index 9) for the forward method, and 24-36% (severity index 7) to 48-57% (severity index 9) for the inverse method (Fig. 4b). Nonetheless, for all the analyzed severity cases, the inverse method shows a lower deviation from realization to realization, as well as a lower dependence to bead density sampling.

**Figure 4:**
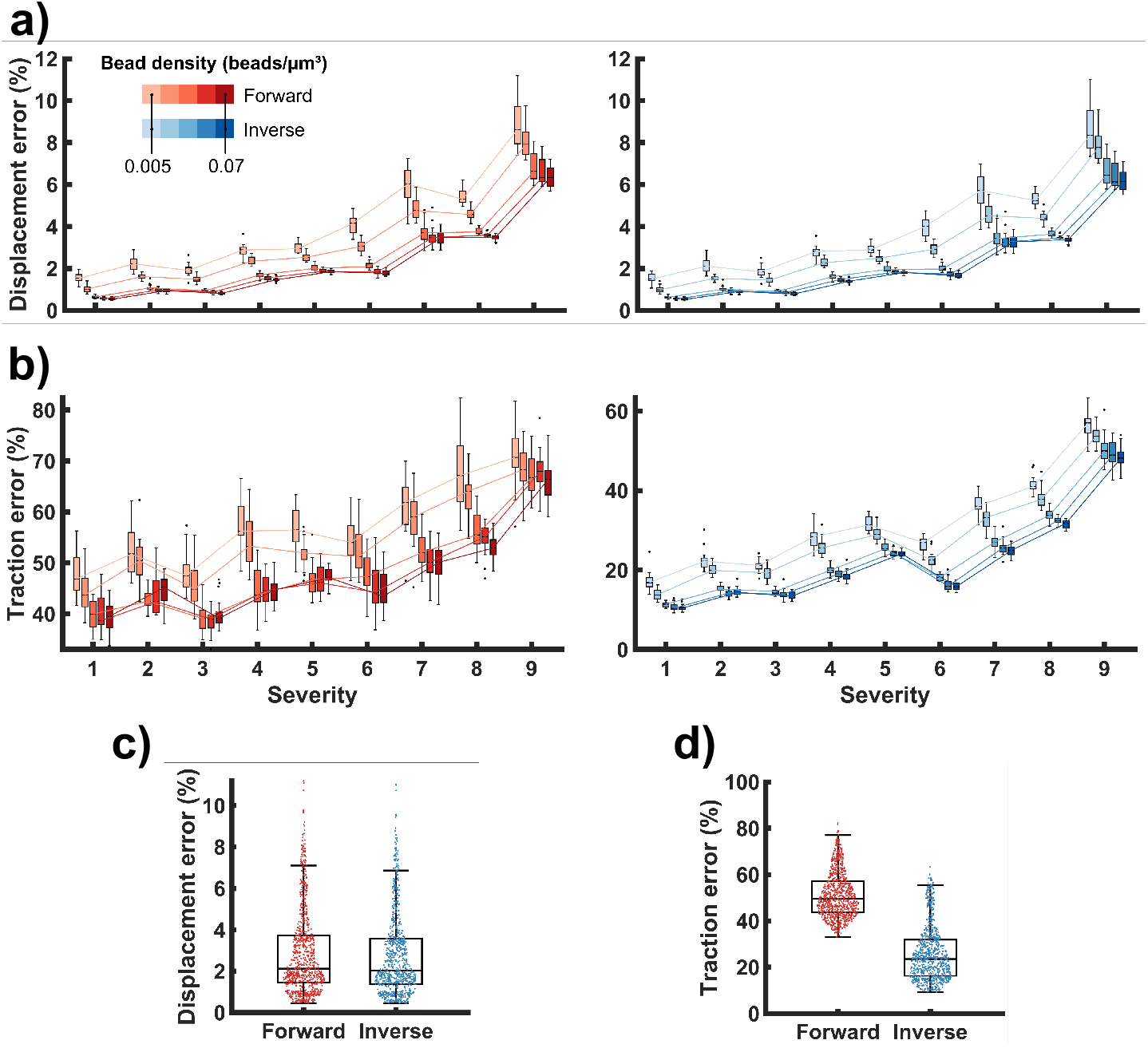
Effect of focal adhesion size and distribution (and associated severity index) and bead density on 3D TFM accuracy. (a) Displacement error with increasing severity using the forward (left) and inverse (right) methods. Boxes are colored with different shades of red (forward) or blue (inverse) according to a bead density from 0.005 to 0.07 beads/*μm*^3^. The median values from each box are connected through straight lines along the different severity values to ease the visualization of the trend. Each box includes the results from 20 bead realizations. (b) Traction error with increasing severity using the forward (left) and inverse (right) methods. Notice the different scales for the vertical axis of the left and right graph. (c) Overall displacement error for the forward and inverse methods. Each box includes the 900 results (depicted as scatter points) from all the analyzed cases (20 realizations *×* 5 bead densities *×* 9 severity cases). (d) Overall traction error for the forward and inverse methods.

A qualitative assessment of the differences between the recovered traction patterns of the two methods is provided in Fig. 5. For low severity (severity 1, Fig. 5a), the *GT* traction pattern is smooth and spread over the four protrusions of the sprout, with the main branch of the sprout not displaying any tractions. The inverse method retrieves similar patterns, with tractions that are mainly located at the protrusions, and the main branch being relatively traction-free. The forward method obtains less precise patterns, in a sense that they are less smooth, with missing tractions in certain parts of the protrusions (see pink arrows in Fig. 5a) and also displaying tractions at the main branch of the sprout (see white arrows in Fig. 5a). Traction peaks are more accurately retrieved by the inverse method (see close-ups of region of interest in Fig. 5a: traction peak around 67 Pa for *GT* and inverse method and around 35 Pa for the forward method). For high severity (severity 9, Fig. 5b), the *GT* traction magnitude pattern is rather sparse and displays areas where tractions decay rapidly to zero. In line with the error quantification in Fig. 4b, the magnitude of these traction peaks is not accurately recovered by either method. Nevertheless, the inverse method provides traction peaks that, while they underestimate the *GT* magnitude, are located in areas closer to the ground truth (see arrows and close-up of the regions of interest in Fig. 5b: traction peak around 250 Pa for *GT*, around 80 Pa for the inverse method and missing peak for the forward method). Further discussion of the results are provided in section 4.

**Figure 5:**
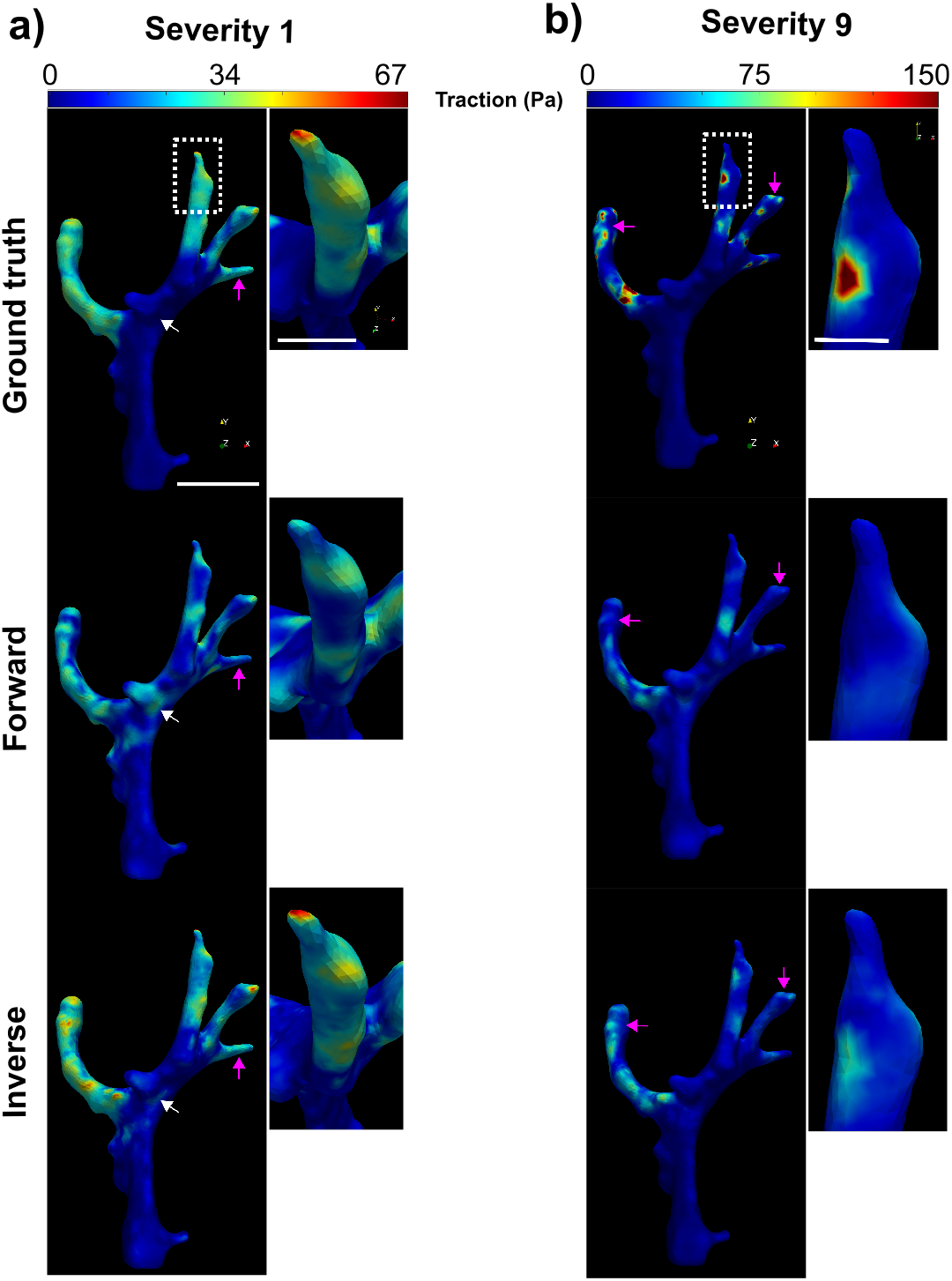
Qualitative representation of the traction patterns of the ground truth (upper row) and the traction recovery by means of the forward (central row) and the inverse (bottom row) methods for one realization of bead density 0.07 beads/*μm*^3^. (a) Traction patterns for the lowest severity (severity 1) for the full sprout (left; scale bar: 21 *μ*m) and for the region of interest depicted with a dotted rectangle (right; scale bar: 7.5 *μ*m). Note that the coordinate system has been rotated for the region of interest to ease the visualization of the traction pattern. Pink arrows indicate areas where the forward method misses tractions and white arrows indicate areas where the forward method displays tractions at the main branch of the sprout. (b) Traction patterns for the highest severity (severity 9) for the full sprout (left; same scale as in (a)) and for the region of interest depicted with a dotted rectangle (right; scale bar: 4.3 *μ*m). Pink arrows indicate other areas where the inverse method captures traction peaks near the *GT*, while the the forward method misses them. Note that the colorbar has been adjusted to ease the visualization of traction patterns in the forward and the inverse method and allow for visual comparison. For the region of interest, the magnitude of the *GT* traction peak shown is around 250 Pa.

### 3.2. 3D TFM of real angiogenic sprouts (in vitro)

Human umbilical vein endothelial cells (HUVECs) were cultured to invade a PEG hydrogel and imaged by means of confocal microscopy (see details in Appendix A). Both the highest and lowest bead densities that were tested in the *in silico* simulations, i.e., 0.07 and 0.005 beads/*μm*^3^, were used. Angiogenic sprouts invaded the 3D PEG hydrogel (elastic modulus of 200Pa and Poisson’s ratio of 0.3[–]) and generated hydrogel displacements that were visualised by means of 200nm fluorescent beads (see Fig. 6a and Fig. 7a). A *relaxed* state of the hydrogel was obtained after inhibiting the cell’s cytoskeletal forces with Cytochalasin D (CytoD). Displacements were measured by means of FFD in a region of interest around the selected sprouts (see 6b and 7b for 0.07 and 0.005 beads/*μm*^3^, respectively). Cellular tractions were recovered using both the forward and the inverse methods. While for experimental cases there is no ground truth to assess accuracy, we can nevertheless compare the tractions retrieved by both methods. The resultant traction maps are shown in Fig. 6c,e and Fig. 7c,e. For the high bead density case (0.07 beads/*μm*^3^; Fig. 6), the forward method retrieved traction patterns of relatively low magnitude (up to 100Pa) and traction peaks that did not necessarily occur near the protrusion tips (Fig. 6c). Moreover, traction vectors did not show clear pulling patterns as they were often not parallel to the protrusion directions (Fig. 6d). In contrast, the inverse method lead to higher traction peaks (150-240Pa) that were mainly located near the protrusion tips (Fig. 6e). Moreover, traction vectors at the protrusion tips were found to be more parallel to the protrusion directions and indicated pulling forces (Fig. 6f), in line with previous reports on displacements and forces around invading sprouts ([48, 56]).

**Figure 6:**
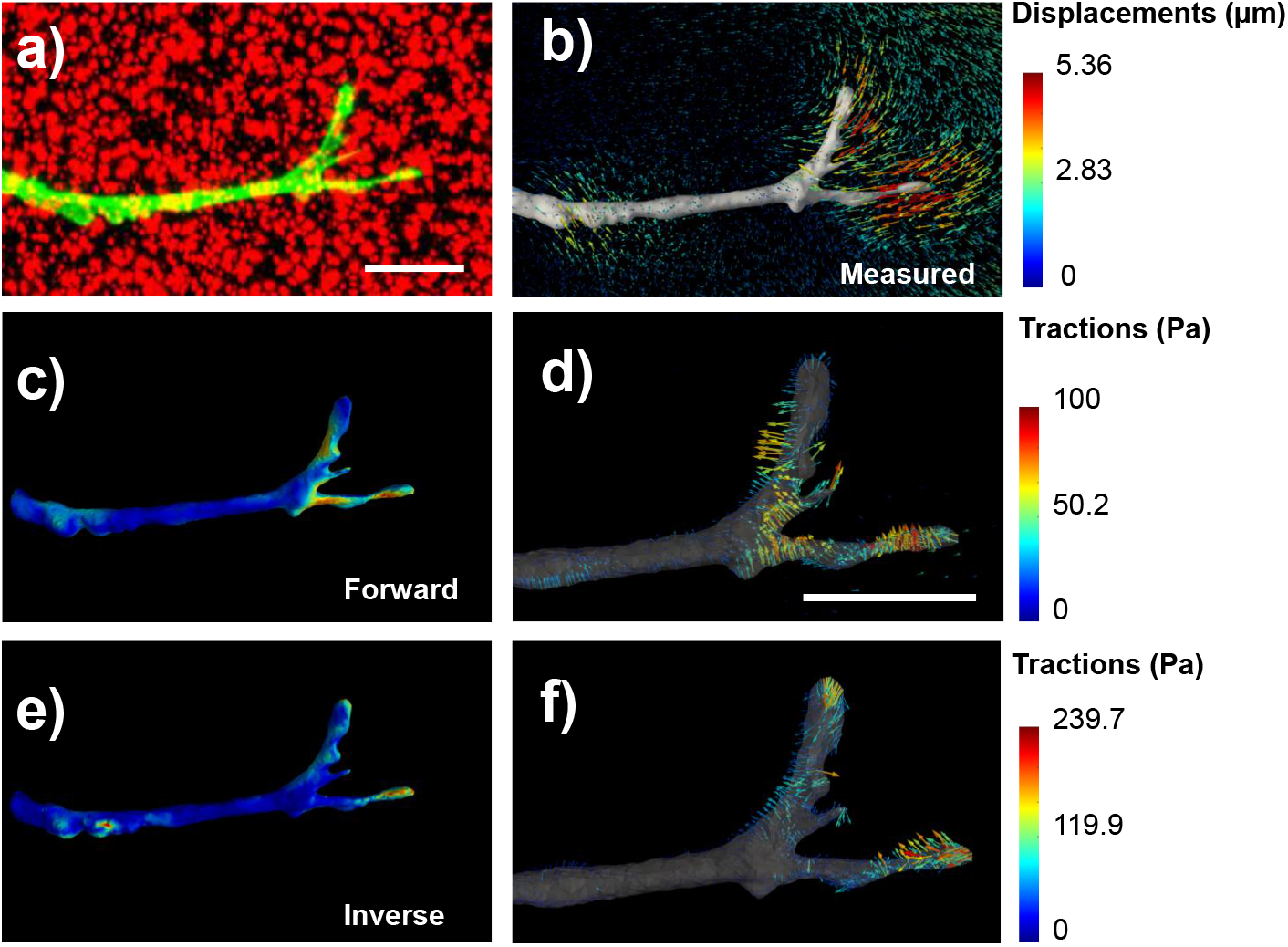
*In vitro* results of 3D TFM of an angiogenic sprout invading a PEG hydrogel that contained 0.07 beads/*μm*^3^. (a) Maximum intensity projection of the image stack (cells in green, beads in red) by means of confocal microscopy. (b) Displacement field measurement by means of FFD. (c) Calculated traction magnitude at the cell surface by means of the forward method. (d) Calculated traction vectors by means of the forward method. (e) Calculated traction magnitude at the cell surface by means of the inverse method. (d) Calculated traction vectors by means of the inverse method. Scale bars: 23 *μ*m

**Figure 7:**
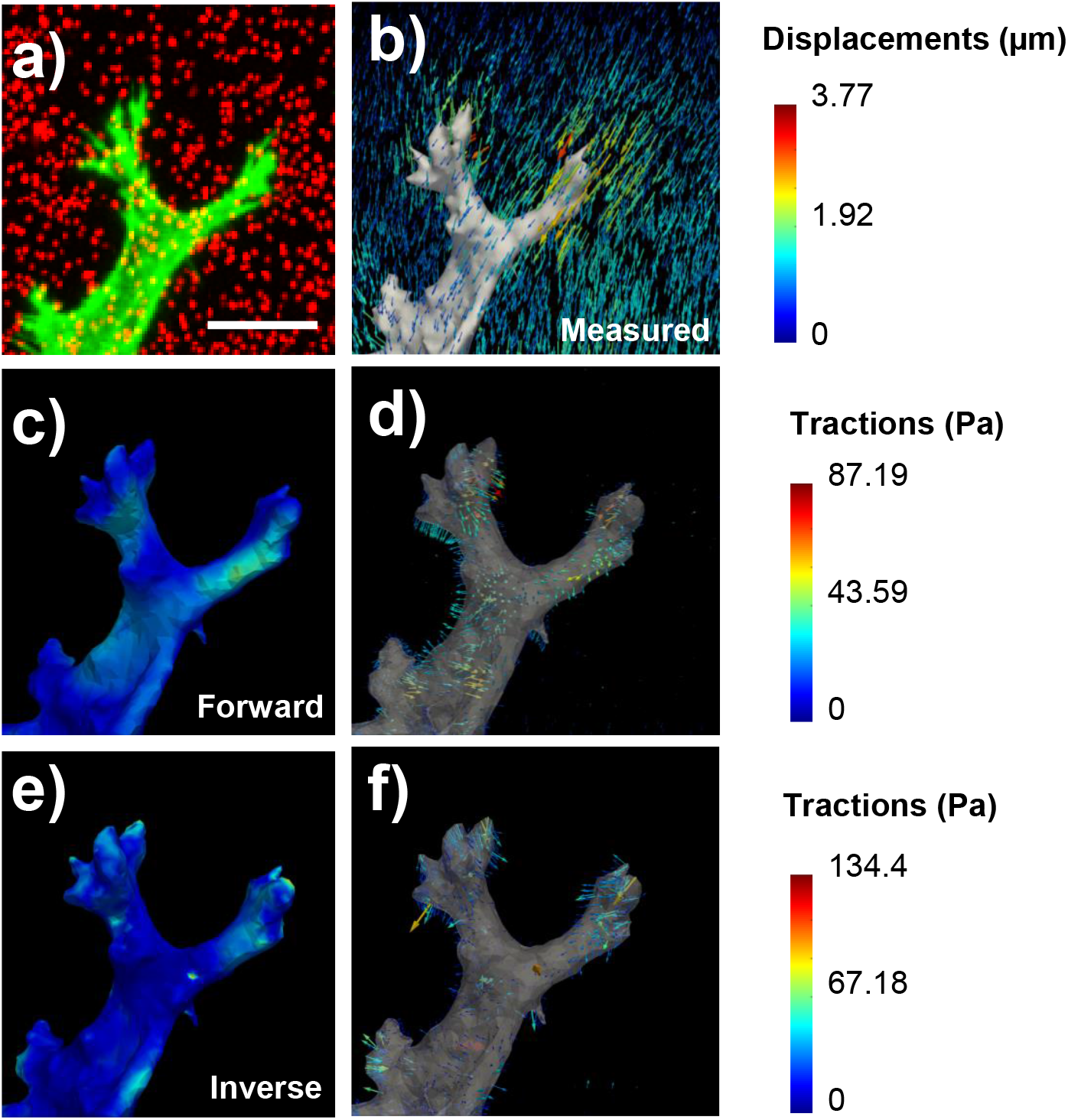
*In vitro* results of 3D TFM of an angiogenic sprout invading a PEG hydrogel that contained 0.005 beads/*μm*^3^. (a) Maximum intensity projection of the image stack (cells in green, beads in red) by means of confocal microscopy. (b) Displacement field measurement by means of FFD. (c) Calculated traction magnitude at the cell surface by means of the forward method. (d) Calculated traction vectors by means of the forward method. (e) Calculated traction magnitude at the cell surface by means of the inverse method. (d) Calculated traction vectors by means of the inverse method. Scale bar: 15 *μ*m

For the low bead density case (0.005 beads/*μm*^3^; Fig. 7), the forward method again retrieves tractions of a lower magnitude compared to the inverse method (90 vs 130 Pa, respectively; Fig. 7c,e). In this case, while the pulling pattern is less pronounced compared to the high bead density case for both methods (Fig. 7d,f), traction peaks recovered by the inverse method are again more localized towards the tips of the protrusions.

## 4. Discussion

3D TFM has the potential to unravel cell mechanical time-dependent interactions with their 3D microenvironment and their role in physiological or pathological processes. However, selecting an adequate traction recovery method that is sufficiently accurate is critical in order to avoid obscuring physiological information (such as, for e.g., differences between relevant conditions) due to traction recovery errors. While the accuracy requirements may differ depending on the application, rigorous and realistic characterization of the expected errors in traction recovery is also crucial for the correct interpretation of experimental results.

In this study, we proposed an advanced validation framework for 3D TFM that includes a real sprout geometry, systematic variation of focal adhesion size and distribution and microscopy bead image simulations. By combining these aspects we established a more realistic and rigorous framework than those encountered in the literature. Within this framework, we thoroughly characterized the accuracy limits of the forward and the inverse traction recovery methods for the considered experimental conditions. We found that 3D TFM accuracy is particularly compromised when calculating cellular tractions associated with small and sparse FAs (in this work, referred to as high severity cases). Small and sparse FAs create small traction areas on the cell’s surface that lead to high gradients in the displacement field of the ECM. The availability of beads in those high gradient areas is reduced, which present more difficulties to sample them properly (see also Appendix F). However, our inverse traction recovery method showed a substantial improvement in accuracy compared to the forward method (errors 2 to 3 times lower) and errors typically below 20% for the highest bead density and most of the severity cases. Moreover, the inverse method provided traction patterns that are closer to the ground truth (similar location of the traction peaks and common traction-free areas). By recovering cellular tractions around angiogenic sprouts in an in vitro model of angiogenesis, the inverse method also showed clearer pulling patterns mainly located near the protrusion tips, further supporting its superiority over the forward method.

This framework is applicable to any 3D TFM experimental setup and can be used to correctly couple the experimental and computational aspects of 3D TFM. While specific experimental characteristics (such as the cell geometry, the microscope PSF or the image voxel size) used in this study could differ from other experiments, the framework can still be used by simply using a different cell geometry, fitting a new PSF and changing the voxel size. Other bead densities outside the range presented here (from 0.005 to 0.07 beads/*μm*^3^) could also be tested. However, researchers must verify that the voxel size is sufficiently small to capture bead displacements and that the bead density is experimentally feasible (avoiding bead clumps in the hydrogel and ensuring that the mechanical properties of the hydrogel and cell behavior are not affected). Moreover, this framework is not limited to hydrogels with linear elastic behavior, since both the forward and our inverse method are compatible with nonlinear elasticity ([51]). Future work could also focus on incorporating knowledge about the location of FAs into the 3D TFM workflow. While we covered a wide range of FA size and distributions in our simulations (see Fig. 2d), FAs may differ depending on the type of cell, adhesiveness and stiffness of the matrix, etc. The inclusion of prior knowledge on the size and distribution of FAs (e.g. by means of FA protein labeling and imaging) in this framework may lead to more specific conclusions for a given experimental application. This could also be used as prior knowledge to further constrain the solution obtained by the inverse method (either the one proposed here or a standard, i.e. Thikonov based linear elastic, inverse method [48]). By applying a physical constraint that regularized the results such that non-zero nodal reaction forces are found only at the cell boundary domain we obtained a 2-fold improvement of traction accuracy with respect to the forward method. The latter method solely relies on the measured displacement field to retrieve tractions. We expect that the inclusion of extra prior knowledge such as the location of FAs could lead to further enhancement of the accuracy of the inverse method, especially in the most challenging (high severity) cases.

## 5. Conclusions

Our study proposes an in silico validation framework for 3D TFM that can be used to select the optimal coupling of experimental (bead density) and computational (traction recovery method) steps. Moreover, it allows for a thorough characterization of the accuracy limits of a traction recovery methods. Our inverse traction recovery method in combination with a high (yet experimentally feasible) bead density provided the most accurate results. By systematically varying FA size and distribution, this framework also revealed the inherent accuracy limitation of 3D TFM, which is linked to small and sparse FAs. Future work should contemplate incorporating prior knowledge on the size, distribution and location of FAs into 3D TFM workflows to further overcome this inherent limitation.

## 6. Acknowledgements

The authors thank Dr Marie-Mo Vaeyens for her valuable input on experimental data acquisition. J. B.-F. was supported by the Research Foundation Flanders (FWO) (travel grant for a long stay abroad to J. B.-F., FWO grant V413019N). J. B.-F., A. R. and H. V. O. were supported by KU Leuven internal funding C14/17/111. A.S. was supported by the FWO SB grant 1S68818N. J. A. S.-H. was supported by the José Castillejo fellowship of the Ministerio de Educación, Cultura y Deporte of Spain, grant number CAS17/00096, and Spanish Ministry of Economy and Competitiveness (MINECO) through the project PGC2018-097257-B-C31. H. V. O. was supported by the European Research Council under the European Union’s Seventh Framework Program (FP7/20072013)/ERC Grant Agreement No. 308223), by an FWO grant (G087018N) and by an FWO/Hercules infrastructure grant (G0H6316N). The financial support is gratefully acknowledged.

## Appendix A. Experimental data acquisition

Angiogenesis is the formation of new blood vessels from existing vasculature ([62, 63]). *In-vitro* models of angiogenesis allow for diffusion of pro-angiogenic signals through a hydrogel that mimics the ECM to promote angiogenic sprouting from a layer of endothelial cells. Leading migratory endothelial cells (tip cells) generate invasive protrusions into the hydrogel and are followed by other endothelial cells (stalk cells) to contribute to the lengthening and ultimate formation of a blood vessel. The angiogenic sprouts used in this study were obtained from a similar experimental assay as the one presented in [56] except that we used a Polyethylene glycol hydrogel (PEG) instead of a collagen hydrogel (see photograph and schematic of the in vitro model in Fig. A.8a,b). Briefly, Human umbilical vein endothelial cells (HUVECs) (Angio-Proteomie, Boston, MA) were cultured in complete endothelial growth medium (EGM-2, Lonza) and used at passage 4. One day before hydrogel preparation, cells were transduced with adenoviral LifeAct-GFP2 (Ibidi). Enzymatically crosslinked poly-ethylene glycol (PEG) hydrogels comprised of an MMP-sensitive peptide modified PEG precursor (8-arm 40kDa), Lys-RGD peptide (Pepmic), and 0.1 *μ*M Sphingosine-1-Phosphate (Sigma-Aldrich) were suspended with 200nm red fluorescent carboxylated polystyrene microspheres (ThermoFischer). The PEG solution was pipetted into the imaging chamber attached to the glass bottom of a petri dish, and further allowed to crosslink for 10 minutes at room temperature. A confluent cell monolayer was then achieved by seeding 50,000 LifeAct transduced HUVECs and incubating the dish vertically for 1 hour at 37 °C, 5% *CO*_2_ to allow cell adhesion onto the PEG meniscus. Finally, EGM-2 was added and dishes were placed horizontally in the incubator for 16 hours before experimentation. The angiogenic sprouts were imaged by means of confocal microscopy using a Leica SP8 with a 25x 0.95 NA water-immersion objective (see example images in Fig. A.8c, d). The green fluorescent cell channel and the red fluorescent beads channel were simultaneously imaged. First, the stressed state was acquired immediately after placing the dish upon the stage and locating the sprout of interest. Second, cells were treated with Cytochalasin D (dissolved in DMSO, Sigma Aldrich) at 4 *μ*M for 50 minutes. Finally, the relaxed state was acquired in the same location.

## Appendix B. Sprout branch segmentation and extraction of principal direction

First, the 3D binary (segmented) image of the sprout was *skeletonized* ([64]) to reduce the cellular surface to single lines or paths without changing its over-all shape. Every skeleton path is identified by junction points (skeleton voxels where different branches meet) and end points (skeleton voxels where branches end).

Second, pairs of junction points and end points were manually selected for the five regions. The sequence of connected voxels with minimum geodesic distance between each pair of points was automatically calculated to get the skeleton path of every region.

Third, each skeleton path corresponding to sprout branches was dilated ([65]) iteratively until its intersection with the binary mask covered the entire branch region. The resultant four binary intersections were used as branch segmentations.

Finally, each skeleton path corresponding to sprout branches was divided into two sub-paths with equal number of voxels and their principal direction vector was computed using principal component analysis (PCA). Therefore, each segmented branch had two assigned direction vectors that were used to determine the direction of the applied tractions in the following simulation steps. The direction of these vectors was corrected to be oriented towards the cell body as experimental data typically show ([39, 35]).

**Figure A.8:**
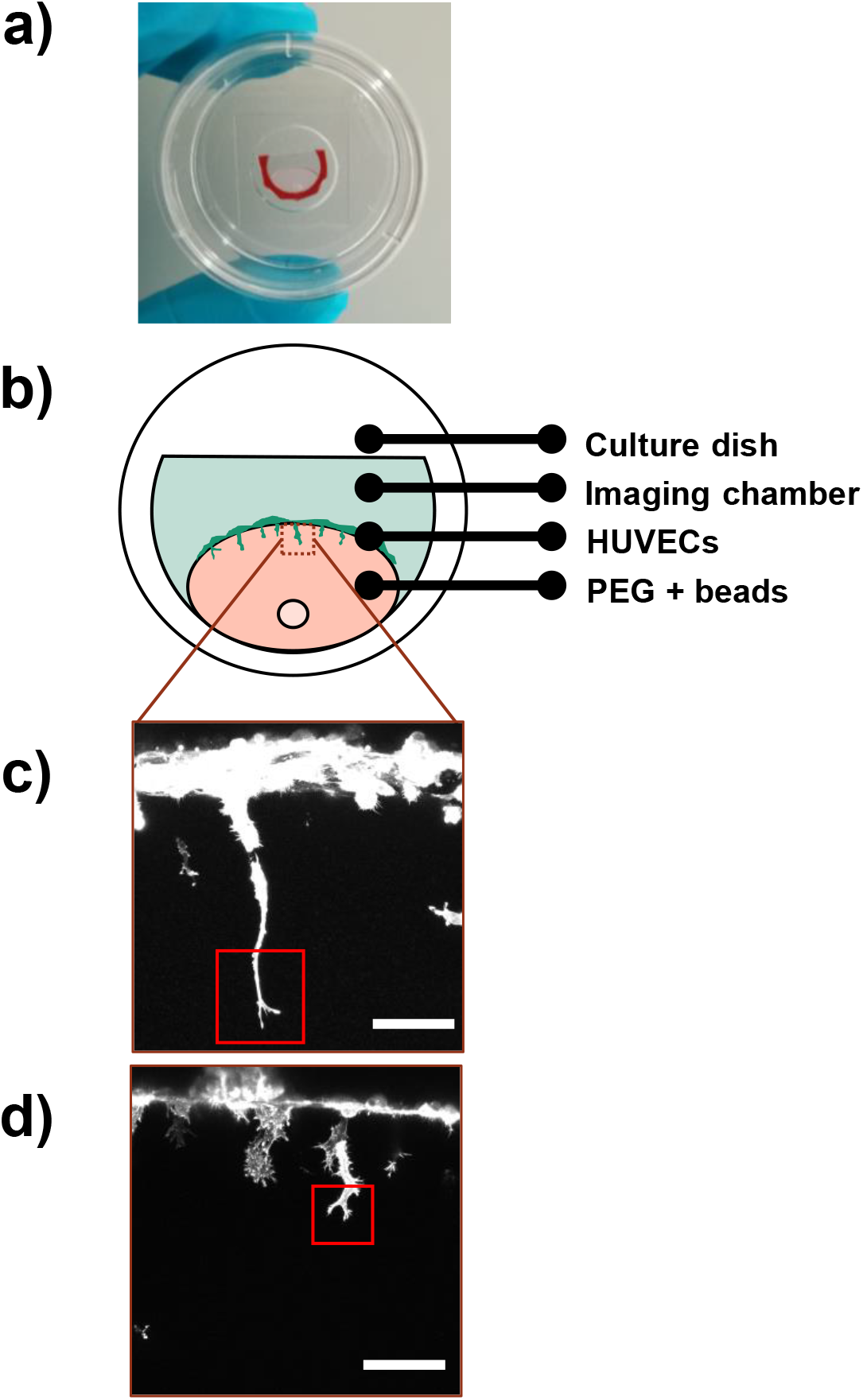
(a) Photograph of the imaging chamber of the in vitro model of angiogenesis used in this study. (b) Schematic of the in vitro model of angiogenesis showing the disposition of the endothelial cell layer and sprouting. The brown dotted square represents a typical imaged region that includes at least one angiogenic sprout. (c, d) Maximum intensity projections of the images acquired by means of confocal microscopy of the sprouts used in a PEG with a bead density of 0.07 and of 0.005 beads/*μm*^3^, respectively. The red rectangles indicate the regions of interest used in Fig. 6. Scale bars: 87 *μ*m.

## Appendix C. Ground truth nodal force simulation and severity calculation

The sum of the magnitude of all the nodal forces was constant for all the FA distributions and fixed at 70nN. In other words, for each FA distribution (see Fig. 2), 70nN of force magnitude was equally distributed over all the focal adhesion nodes. Table C.1 shows the total number of FA nodes (of all four protrusions) and the magnitude of the force applied in each node, for every case of study. It can be observed that while the total force is constant, different nodal force magnitudes are applied for each case, depending on the number of FA nodes. This, combined with the spatial distribution of the focal adhesion nodes, leads to differences in traction field heterogeneity (non-smoothness). We defined a severity index (from 1 to 9) that describes the heterogeneity of the traction field and that was associated to the coefficient of variation (COV) according to the formula: 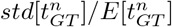, where 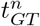 is the value of the magnitude of the ground truth traction at the *n* nodes of the sprout boundary domain and std[] and E[] stand for standard deviation and mean respectively. We computed this metric for each of the 9 cases defined by combining different focal adhesion distributions and sizes (see Fig. C.9 and section 2.1) and assigned them a severity index from 1 to 9 (as shown in Fig. 2).

## Appendix D. Point-spread function simulation

In this study, a point spread function (PSF) was generated by fitting a 3D Gaussian profile ([66, 55]) to randomly distributed 200nm beads within the volume of experimental confocal microscopy images. Briefly, each bead affected by the PSF is expressed as a Gaussian blob as:

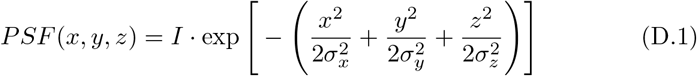

where *I* is the bead maximum intensity value and *σ* = (*σ*_*x*_, *σ*_*y*_, *σ*_*z*_) is the 3D standard deviation. *I* was set to randomly vary within the intensity range [240,255] to simulate differences in bead fluorescence emissions and parameter (*σ*_*x*_, *σ*_*y*_, *σ*_*z*_) = ([1, 1, 1.58]) was fixed based on the fitting to experimental data. Gaussian blobs were simulated using function `gaussianblob` from the DipImage library [67]. The spatial resolution or voxel size of the images was set to match that of the experimental setup: 0.35 *μm* in the XY-plane and 1 *μm* in the Z-axis.

**Figure C.9:**
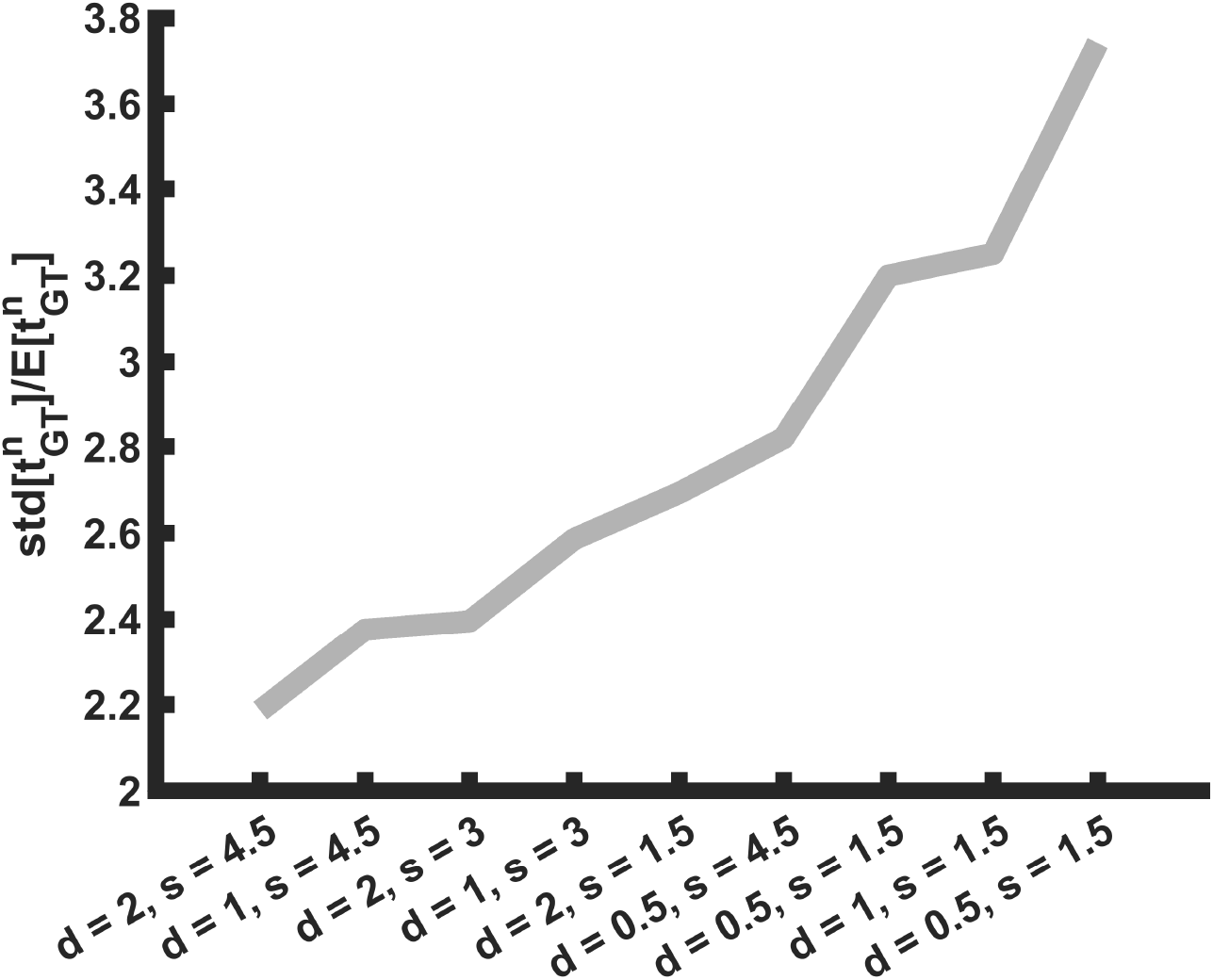
Calculation of the severity index for each of the ground truth cases. d stands for density (in %) and s stands for size (in *μ*m) (see also section 2.1).

**Table C.1:** Number of FA nodes for each severity case and magnitude of the applied nodal forces.

| Severity case | Number of FA nodes | Nodal force magnitude (pN) |
| --- | --- | --- |
| 1 | 2764 | 25.33 |
| 2 | 1855 | 37.74 |
| 3 | 1485 | 47.14 |
| 4 | 855 | 81.87 |
| 5 | 284 | 246.48 |
| 6 | 1088 | 64.34 |
| 7 | 469 | 149.25 |
| 8 | 142 | 492.96 |
| 9 | 71 | 985.92 |

In this study, 5 bead densities were considered, namely, 0.005, 0.01, 0.03, 0.05 and 0.07 beads/*μm*^3^. Since bead density can be found in literature in different units we provide Table D.2 to ease the comparison with other works.

## Appendix E. Displacement calculation by means of FFD

Briefly, FFD overlays a regular mesh to a *stressed* image (i.e. deformed hydrogel) and identifies a mesh transformation model that wraps it to match a *relaxed* image (i.e. undeformed hydrogel). The positions of nodes of the mesh, which are the control points of the multivariate B-spline functions, are iteratively tuned to compute an optimal transformation. In this work, image registration parameters used in [68] were selected, namely, normalized correlation coefficient as the similarity metric, gradient descent method with adaptive estimation of the step size as the optimization strategy and three-level coarse-to-fine multiscaling technique was applied. After a sensitivity analysis, the distance between control points was set to 3 microns for all the simulations. Registration was successful for all the cases, showing high registration metric results (around 98-99%).

**Table D.2:** Bead densities used in this study and its equivalences in other units (namely, beads per image voxel in % and beads per mL of hydrogel) to ease their comparison with other works.

| Bead density (beads/$\mu m^3$) | Beads per voxel (%) | Beads per mL ($\cdot 10^9$) |
| --- | --- | --- |
| 0.005 | 0.06 | 5 |
| 0.01 | 0.12 | 10 |
| 0.03 | 0.37 | 30 |
| 0.05 | 0.61 | 50 |
| 0.07 | 0.86 | 70 |

## Appendix F. Bead density and sampling close to focal adhesions

Small and sparse FAs create small traction areas on the cell’s surface that lead to high gradients in the displacement field of the ECM and that present more difficulties to sample them properly. To illustrate that the availability of beads within high gradient areas is reduced, we obtained the bead positions of 3 realizations from our in silico simulations for bead densities 0.005 and 0.07 beads/*μm*^3^ and for severity cases 1 and 9. Fig.F.10a,b shows the major principal stress in the ECM, which is a function of the gradient of the displacement field, for *GT* severity cases 1 and 9. When a low bead density is used (0.005 beads/*μm*^3^) beads rarely fall within the higher stress areas (for both severity cases). When a high bead density is used (0.07 beads/*μm*^3^), more beads fall within higher stress areas for the severity case 1 (Fig. F.10a; lower row). However, it can be seen that some high stress areas are not sampled. The same is true for severity case 9: since the higher stress areas are smaller and sparser there are few or no beads that can be used as sampling points of higher stress (high displacement gradient) areas, even for the high bead density (Fig. F.10b, lower row).

This illustrates the inherent problem of 3D TFM: an increase of the bead density only leads to a modest improvement of the sampling of high stress areas (close vicinity of FAs) in the ECM. For a given distribution of FAs (see example for severity case 5 in Fig. F.10c), an increase of the bead density from 0.005 to 0.07 beads/*μm*^3^ rapidly increases the number of beads away from the FAs: the number of beads within a distance of 1-10 *μm* from the FAs increases from around 100 to more than 1000 (proportional to the considered volume around the FA, and therefore proportional to the third power of the distance to the FA). However, closer to the FAs the number of beads increases at a much slower rate (following from the much smaller volume to be sampled): the number of beads within a distance of 0-1 *μm* from the FAs remains below 10. Given an increase of the bead density in steps of 0.02 beads/*μm*^3^, the number of beads within a distance of 0.5, 1, 5 and 10 *μm* from the FAs increases on average with a factor 1.67, 8, 369.7 and 1209 respectively (see Fig. F.10d).

Please note that the plots shown in Fig. F.10a,b are different from the one shown in Fig. 3b. Here, only the bead centroids in the central slice are depicted while Fig. 3b shows the central slice of a simulated bead image, where due to the PSF, one can see parts of beads located in adjacent planes. Moreover, the size of the scatter points used for these plots is larger than a pixel, to ease their visualization.

## Appendix G. Angle deviation in traction recovery

The angle deviation between the tractions recovered by the forward and the inverse method was measured using the following formula:

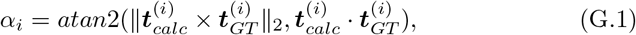

where *atan*2 is the four-quadrant inverse tangent, *|| • ||*_2_ is the L2-norm, *×* is the cross product and *·* is the dot product. An angle of 0° indicates a perfect alignment between the calculated traction 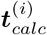 and the ground truth traction 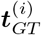 at node *i*. An angle of 180° indicates parallel vectors pointing in opposite direction. For each analyzed case (9 realizations *×* 5 bead densities *×* 9 severity cases), we averaged the angles at the nodes where the magnitude of the *GT* traction field was above the threshold: 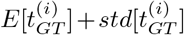. Therefore, we obtained one average angle for each case. The results are shown in Fig. G.11. The inverse method retrieves traction vectors that are more aligned with the ground truth than the forward method (20-30° vs 40-50° of deviation for most of the cases, respectively). Again, high severity cases lead to higher deviations (up to 40° for the inverse method and up to 70° for the forward method).

**Figure F.10:**
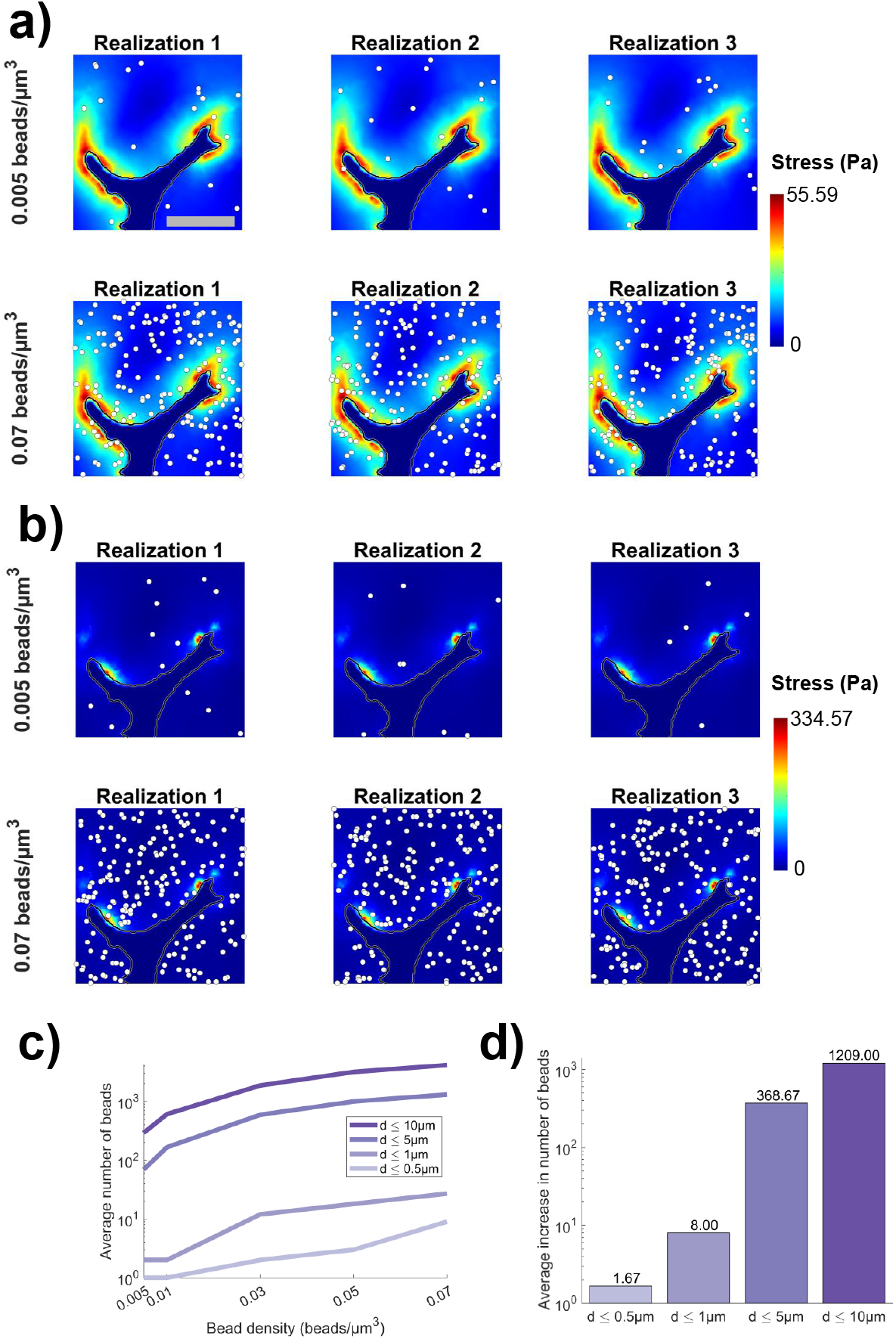
(a) Central xy plane of the *GT* major principal stress (colormap) and sprout boundary (in black) for severity case 1 and bead realizations of 0.005 (upper row) and 0.07 (lower row) beads/*μm*^3^. Three bead realizations are shown. The centroid of each bead present in the central xy plane is depicted with white scatter points. Scale bar: 22*μm*. (b) Same as in (a), but for severity case 9. (c) Average number of beads (over 20 bead realizations) present within a distance ‘d’ (measured from bead centroid to FA node) from an FA as a function of bead density for severity case 5. (d) Average (over the ranges [0.01,0.03], [0.03,0.05] and [0.05,0.07] beads/*μm*^3^) slopes of the curves in (c).

**Figure G.11:**
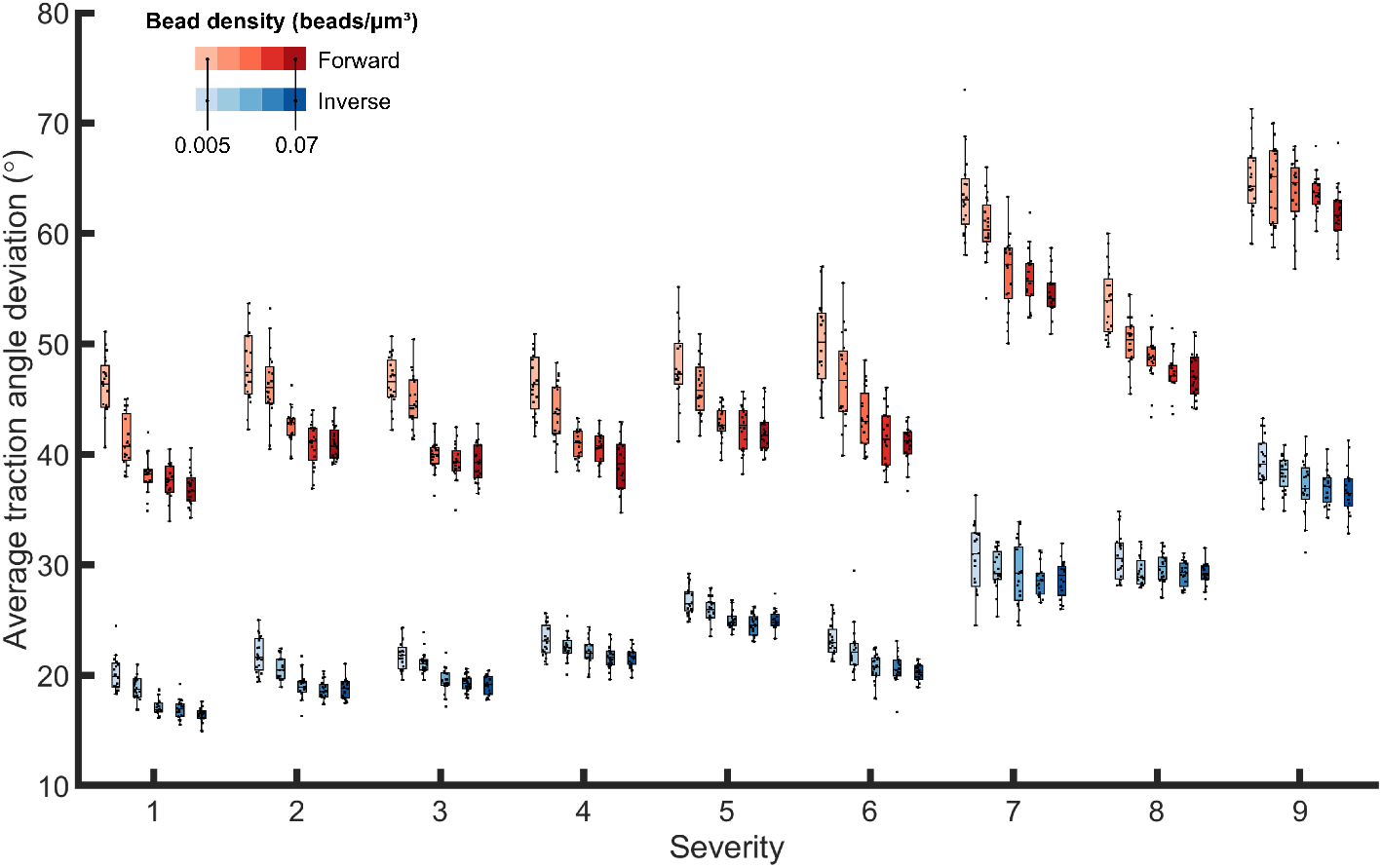
Average traction angle deviation with increasing severity. Boxes are colored with different shades of red (forward) or blue (inverse) according to a bead density from 0.005 to 0.07 beads/*μm*^3^).

## Notes

### Competing Interest Statement

The authors have declared no competing interest.

## References

[1] D. Ingber, Mechanobiology and diseases of mechanotransduction, Annals of Medicine 35 (8) (2003) 564–577. doi:10.1080/07853890310016333. URL http://www.tandfonline.com/doi/full/10.1080/07853890310016333

[2] M. A. Wozniak, C. S. Chen, Mechanotransduction in development: a growing role for contractility., Nature reviews. Molecular cell biology 10 (1) (2009) 34–43. doi:10.1038/nrm2592. URL http://www.ncbi.nlm.nih.gov/pubmed/19197330 http://www.pubmedcentral.nih.gov/articlerender.fcgi?artid=PMC2952188

[3] F. Guilak, D. M. Cohen, B. T. Estes, J. M. Gimble, W. Liedtke, C. S. Chen, Control of Stem Cell Fate by Physical Interactions with the Extracellular Matrix, Cell Stem Cell 5 (1) (2009) 17–26. doi:10.1016/J.STEM.2009.06.016. URL https://www.cell.com/cell-stem-cell/fulltext/S1934-5909{%}2809{%}2900293-8

[4] S. Kumar, V. M. Weaver, Mechanics, malignancy, and metastasis: The force journey of a tumor cell, Cancer metastasis reviews 28 (1-2) (2009) 113. doi:10.1007/S10555-008-9173-4. URL https://www.ncbi.nlm.nih.gov/pmc/articles/PMC2658728/

[5] M. Ehrbar, A. Sala, P. Lienemann, A. Ranga, K. Mosiewicz, A. Bittermann, S. C. Rizzi, F. E. Weber, M. P. Lutolf, Elucidating the role of matrix stiffness in 3D cell migration and remodeling, Biophysical Journal 100 (2) (2011) 284–293. doi:10.1016/j.bpj.2010.11.082.

[6] J. Zhong, Y. Yang, L. Liao, C. Zhang, Matrix stiffness-regulated cellular functions under different dimensionalities, Biomaterials science 8 (10) (2020) 2734–2755. doi:10.1039/c9bm01809c. URL https://pubs.rsc.org/en/content/articlelanding/2020/bm/c9bm01809c

[7] D. E. Discher, P. Janmey, Y.-L. Wang, Tissue cells feel and respond to the stiffness of their substrate., Science (New York, N.Y.) 310 (5751) (2005) 1139–43. doi:10.1126/science.1116995. URL http://www.ncbi.nlm.nih.gov/pubmed/16293750

[8] A. J. Engler, S. Sen, H. L. Sweeney, D. E. Discher, Matrix Elasticity Directs Stem Cell Lineage Specification, Cell 126 (4) (2006) 677–689. doi:10.1016/J.CELL.2006.06.044. URL https://www.sciencedirect.com/science/article/pii/S0092867406009615

[9] N. Huebsch, P. R. Arany, A. S. Mao, D. Shvartsman, O. A. Ali, S. A. Bencherif, J. Rivera-Feliciano, D. J. Mooney, Harnessing traction-mediated manipulation of the cell/matrix interface to control stem-cell fate, Nature Materials 9 (6) (2010) 518–526. doi:10.1038/nmat2732. URL http://www.nature.com/articles/nmat2732

[10] L. Wong, A. Kumar, B. Gabela-Zuniga, J. Chua, G. Singh, C. L. Happe, A. J. Engler, Y. Fan, K. E. McCloskey, Substrate stiffness directs diverging vascular fates, Acta Biomaterialia 96 (2019) 321–329. doi:10.1016/j.actbio.2019.07.030.

[11] C. Puckert, E. Tomaskovic-Crook, S. Gambhir, G. G. Wallace, J. M. Crook, M. J. Higgins, Molecular interactions and forces of adhesion between single human neural stem cells and gelatin methacrylate hydrogels of varying stiffness, Acta Biomaterialia 106 (2020) 156–169. doi:10.1016/j.actbio.2020.02.023.

[12] O. Chaudhuri, L. Gu, D. Klumpers, M. Darnell, S. A. Bencherif, J. C. Weaver, N. Huebsch, H.-P. Lee, E. Lippens, G. N. Duda, D. J. Mooney, Hydrogels with tunable stress relaxation regulate stem cell fate and activity., Nature materials 15 (3) (2016) 326–34. doi:10.1038/nmat4489. URL http://www.ncbi.nlm.nih.gov/pubmed/26618884 http://www.pubmedcentral.nih.gov/articlerender.fcgi?artid=PMC4767627

[13] C. N. Salinas, K. S. Anseth, The enhancement of chondrogenic differentiation of human mesenchymal stem cells by enzymatically regulated RGD functionalities, Biomaterials 29 (15) (2008) 2370–2377. doi:10.1016/j.biomaterials.2008.01.035.

[14] J. D. Shields, M. S. Emmett, D. B. Dunn, K. D. Joory, L. M. Sage, H. Rigby, P. S. Mortimer, A. Orlando, J. R. Levick, D. O. Bates, Chemokine-mediated migration of melanoma cells towards lymphatics - A mechanism contributing to metastasis, Oncogene 26 (21) (2007) 2997–3005. doi:10.1038/sj.onc.1210114. URL https://www.nature.com/articles/1210114

[15] K. V. Nguyen-Ngoc, K. J. Cheung, A. Brenot, E. R. Shamir, R. S. Gray, W. C. Hines, P. Yaswen, Z. Werb, A. J. Ewald, ECM microenvironment regulates collective migration and local dissemination in normal and malignant mammary epithelium, Proceedings of the National Academy of Sciences of the United States of America 109 (39) (2012) E2595–E2604. doi:10.1073/pnas.1212834109. URL https://www.pnas.org/content/109/39/E2595 https://www.pnas.org/content/109/39/E2595.abstract

[16] S. H. Medina, B. Bush, M. Cam, E. Sevcik, F. W. DelRio, K. Nandy, J. P. Schneider, Identification of a mechanogenetic link between substrate stiffness and chemotherapeutic response in breast cancer, Biomaterials 202 (2019) 1–11. doi:10.1016/j.biomaterials.2019.02.018.

[17] T. Jiang, J. Zhao, S. Yu, Z. Mao, C. Gao, Y. Zhu, C. Mao, L. Zheng, Untangling the response of bone tumor cells and bone forming cells to matrix stiffness and adhesion ligand density by means of hydrogels, Biomaterials 188 (2019) 130–143. doi:10.1016/j.biomaterials.2018.10.015.

[18] Y. Peng, Z. Chen, Y. Chen, S. Li, Y. Jiang, H. Yang, C. Wu, F. You, C. Zheng, J. Zhu, Y. Tan, X. Qin, Y. Liu, ROCK isoforms differentially modulate cancer cell motility by mechanosensing the substrate stiffness, Acta Biomaterialia 88 (2019) 86–101. doi:10.1016/j.actbio.2019.02.015.

[19] A. Khang, A. Gonzalez Rodriguez, M. E. Schroeder, J. Sansom, E. Lejeune, K. S. Anseth, M. S. Sacks, Quantifying heart valve interstitial cell contractile state using highly tunable poly(ethylene glycol) hydrogels, Acta Biomaterialia 96 (2019) 354–367. doi:10.1016/j.actbio.2019.07.010.

[20] O. Chaudhuri, L. Gu, M. Darnell, D. Klumpers, S. A. Bencherif, J. C. Weaver, N. Huebsch, D. J. Mooney, Substrate stress relaxation regulates cell spreading, Nature Communications 6 (1) (2015) 1–7. doi:10.1038/ncomms7365. URL https://www.nature.com/articles/ncomms7365 https://www.nature.com/articles/ncomms7365/

[21] B. M. Baker, B. Trappmann, W. Y. Wang, M. S. Sakar, I. L. Kim, V. B. Shenoy, J. A. Burdick, C. S. Chen, Cell-mediated fibre recruitment drives extracellular matrix mechanosensing in engineered fibrillar microenvironments, Nature Materials 14 (12) (2015) 1262–1268. doi: 10.1038/nmat4444. URL http://www.nature.com/articles/nmat4444

[22] J. P. Butler, I. M. Tolić-Nørrelykke, B. Fabry, J. J. Fredberg, Traction fields, moments, and strain energy that cells exert on their surroundings, American Journal of Physiology - Cell Physiology 282 (3). URL http://ajpcell.physiology.org/content/282/3/C595

[23] B. Sabass, M. L. Gardel, C. M. Waterman, U. S. Schwarz, High resolution traction force microscopy based on experimental and computational advances., Biophysical journal 94 (1) (2008) 207–20. doi:10.1529/biophysj.107.113670. URL http://www.ncbi.nlm.nih.gov/pubmed/17827246 http://www.pubmedcentral.nih.gov/articlerender.fcgi?artid=PMC2134850

[24] B. M. Baker, C. S. Chen, Deconstructing the third dimension-how 3D culture microenvironments alter cellular cues (jul 2012). doi:10.1242/jcs.079509.

[25] J. Stricker, Y. Aratyn-Schaus, P. W. Oakes, M. L. Gardel, Spatiotemporal Constraints on the Force-Dependent Growth of Focal Adhesions, Biophysical Journal 100 (12) (2011) 2883–2893. doi:10.1016/J.BPJ.2011.05.023. URL https://www.sciencedirect.com/science/article/pii/S0006349511005935?via%3Dihub

[26] C. S. Chen, Mechanotransduction - a field pulling together?, Journal of cell science 121 (Pt 20) (2008) 3285–92. doi:10.1242/jcs.023507. URL http://www.ncbi.nlm.nih.gov/pubmed/18843115

[27] S. I. Fraley, Y. Feng, R. Krishnamurthy, D.-H. Kim, A. Celedon, G. D. Longmore, D. Wirtz, A distinctive role for focal adhesion proteins in three-dimensional cell motility, Nature Cell Biology 12 (6) (2010) 598–604. doi: 10.1038/ncb2062. URL http://www.nature.com/articles/ncb2062

[28] S. I. Fraley, Y. Feng, D. Wirtz, G. D. Longmore, Reply: Reducing background fluorescence reveals adhesions in 3D matrices (jan 2011). doi: 10.1038/ncb0111-5. URL https://www.nature.com/articles/ncb0111-5

[29] K. E. Kubow, A. R. Horwitz, Reducing background fluorescence reveals adhesions in 3D matrices, Nature Cell Biology 13 (1) (2011) 3–5. doi: 10.1038/ncb0111-3. URL http://www.nature.com/doifinder/10.1038/ncb0111-3

[30] K. E. Kubow, S. K. Conrad, A. R. Horwitz, Matrix microarchitecture and myosin II determine adhesion in 3D matrices, Current Biology 23 (17) (2013) 1607–1619. doi:10.1016/j.cub.2013.06.053.

[31] C.-L. Chiu, J. S. Aguilar, C. Y. Tsai, G. Wu, E. Gratton, M. A. Digman, Nanoimaging of Focal Adhesion Dynamics in 3D, PLoS ONE 9 (6) (2014) e99896. doi:10.1371/journal.pone.0099896. URL https://dx.plos.org/10.1371/journal.pone.0099896

[32] J. A. Broussard, N. L. Diggins, S. Hummel, W. Georgescu, V. Quaranta, D. J. Webb, Automated analysis of cell-matrix adhesions in 2D and 3D environments, Scientific Reports 5 (1) (2015) 8124. doi:10.1038/srep08124. URL https://www.nature.com/articles/srep08124

[33] A. D. Doyle, K. M. Yamada, Mechanosensing via cell-matrix adhesions in 3D microenvironments, Experimental Cell Research 343 (1) (2016) 60–66. doi:10.1016/j.yexcr.2015.10.033.

[34] H. Colin-York, D. Shrestha, J. H. Felce, D. Waithe, E. Moeendarbary, S. J. Davis, C. Eggeling, M. Fritzsche, Super-Resolved Traction Force Microscopy (STFM), Nano Letters 16 (4) (2016) 2633–2638. doi:10.1021/acs.nanolett.6b00273. URL http://pubs.acs.org/doi/10.1021/acs.nanolett.6b00273

[35] W. R. Legant, J. S. Miller, B. L. Blakely, D. M. Cohen, G. M. Genin, C. S. Chen, Measurement of mechanical tractions exerted by cells in three-dimensional matrices, Nature Methods 7 (12) (2010) 969–971. doi:10.1038/nmeth.1531. URL http://www.nature.com/articles/nmeth.1531

[36] S. J. Han, Y. Oak, A. Groisman, G. Danuser, Traction microscopy to identify force modulation in subresolution adhesions, Nature Methods 12 (7) (2015) 653–656. doi:10.1038/nmeth.3430. URL https://www.nature.com/articles/nmeth.3430.pdf

[37] I. M. Tolić-Nørrelykke, J. P. Butler, J. Chen, N. Wang, Spatial and temporal traction response in human airway smooth muscle cells, American Journal of Physiology-Cell Physiology 283 (4) (2002) C1254–C1266. doi:10.1152/ajpcell.00169.2002. URL http://www.physiology.org/doi/10.1152/ajpcell.00169.2002

[38] Y. Du, S. C. B. Herath, Q.-g. Wang, D.-a. Wang, H. H. Asada, P. C. Y. Chen, Three-Dimensional Characterization of Mechanical Interactions between Endothelial Cells and Extracellular Matrix during Angiogenic Sprouting, Scientific Reports 6 (1) (2016) 21362. doi:10.1038/srep21362. URL http://www.nature.com/articles/srep21362

[39] A. Jorge-Peñas, A. Izquierdo-Alvarez, R. Aguilar-Cuenca, M. Vicente-Manzanares, J. M. Garcia-Aznar, H. Van Oosterwyck, E. M. De-Juan-Pardo, C. Ortiz-De-Solorzano, A. Muñoz-Barrutia, Free form deformation-based image registration improves accuracy of traction force microscopy, PLoS ONE 10 (12) (2015) 1–22. doi:10.1371/journal.pone.0144184.

[40] C. Franck, S. A. Maskarinec, D. A. Tirrell, G. Ravichandran, G. Genin, Three-Dimensional Traction Force Microscopy: A New Tool for Quantifying Cell-Matrix Interactions, PLoS ONE 6 (3) (2011) e17833. doi: 10.1371/journal.pone.0017833. URL http://dx.plos.org/10.1371/journal.pone.0017833

[41] J. Toyjanova, E. Bar-Kochba, C. López-Fagundo, J. Reichner, D. Hoffman-Kim, C. Franck, High resolution, large deformation 3D traction force microscopy, PLoS ONE 9 (4) (2014) 1–12. doi:10.1371/journal.pone.0090976.

[42] N. Gjorevski, A. S. Piotrowski, V. D. Varner, C. M. Nelson, Dynamic tensile forces drive collective cell migration through three-dimensional extracellular matrices, Scientific reports 5 (2015) 11458. doi:10.1038/srep11458.

[43] S. S. Hur, Y. Zhao, Y.-S. Li, E. Botvinick, S. Chien, Live Cells Exert 3-Dimensional Traction Forces on Their Substrata, Cellular and Molecular Bioengineering 2 (3) (2009) 425–436. doi:10.1007/s12195-009-0082-6. URL http://www.ncbi.nlm.nih.gov/pubmed/19779633 http://www.pubmedcentral.nih.gov/articlerender.fcgi?artid=PMC2749171 http://link.springer.com/10.1007/s12195-009-0082-6

[44] A. S. Piotrowski, V. D. Varner, N. Gjorevski, C. M. Nelson, Three-dimensional traction force microscopy of engineered epithelial tissues, Methods in Molecular Biology 1189 (2015) 191–206. doi:10.1007/978-1-4939-1164-6_13. URL https://link.springer.com/protocol/10.1007/978-1-4939-1164-6{_}13

[45] C. N. Holenstein, C. R. Lendi, N. Wili, J. G. Snedeker, Simulation and evaluation of 3D traction force microscopy, Computer Methods in Biomechanics and Biomedical Engineering 22 (8) (2019) 853–860. doi:10.1080/10255842.2019.1599866. URL https://www.tandfonline.com/doi/full/10.1080/10255842.2019.1599866

[46] U. S. Schwarz, J. R. D. Soiné, Traction force microscopy on soft elastic substrates: A guide to recent computational advances, Biochimica et Biophysica Acta - Molecular Cell Research 1853 (11) (2015) 3095–3104. arXiv:1506.02394v1, doi:10.1016/j.bbamcr.2015.05.028. URL http://dx.doi.org/10.1016/j.bbamcr.2015.05.028

[47] J. Steinwachs, C. Metzner, K. Skodzek, N. Lang, I. Thievessen, C. Mark, S. Münster, K. E. Aifantis, B. Fabry, Three-dimensional force microscopy of cells in biopolymer networks, Nature Methods 13 (2) (2016) 171–176. doi:10.1038/nmeth.3685.

[48] Y. Du, S. Herath, Q. Wang, H. Asada, P. Chen, Determination of Green’s function for three-dimensional traction force reconstruction based on geometry and boundary conditions of cell culture matrices, Acta Biomaterialia 67 (2018) 215–228. doi:10.1016/J.ACTBIO.2017.12.002. URL https://www.sciencedirect.com/science/article/pii/S1742706117307584{#}s0065

[49] D. Song, N. Hugenberg, A. A. Oberai, Three-dimensional traction microscopy with a fiber-based constitutive model, Computer Methods in Applied Mechanics and Engineering 357 (2019) 112579. doi:10.1016/J.CMA.2019.112579. URL https://www.sciencedirect.com/science/article/pii/S004578251930444X

[50] M. Cóndor, J. García-Aznar, An iterative finite element-based method for solving inverse problems in traction force microscopy, Computer Methods and Programs in Biomedicine 182 (2019) 105056. doi:10.1016/j.cmpb.2019.105056. URL https://linkinghub.elsevier.com/retrieve/pii/S0169260719305358

[51] J. A. Sanz-Herrera, J. Barrasa Fano, M. Cóndor, H. Van Oosterwyck, Inverse method based on 3D nonlinear physically constrained minimisation in the framework of traction force microscopy., Soft Matter (in press). doi:10.1039/D0SM00789G. URL http://pubs.rsc.org/en/Content/ArticleLanding/2020/SM/D0SM00789G

[52] C. N. Holenstein, C. R. Lendi, N. Wili, J. G. Snedeker, Simulation and evaluation of 3D traction force microscopy, Computer Methods in Biomechanics and Biomedical Engineering 22 (8) (2019) 853–860. doi: 10.1080/10255842.2019.1599866. URL https://doi.org/10.1080/10255842.2019.1599866

[53] G. Vitale, L. Preziosi, D. Ambrosi, A numerical method for the inverse problem of cell traction in 3D, Inverse Problems 28 (09501317pp). doi: 10.1088/0266-5611/28/9/095013. URL http://iopscience.iop.org/article/10.1088/0266-5611/28/9/095013/pdf

[54] L. Dong, A. A. Oberai, Recovery of cellular traction in three-dimensional nonlinear hyperelastic matrices, Computer Methods in Applied Mechanics and Engineering 314 (2017) 296–313. doi:10.1016/J.CMA.2016.05.020. URL https://www.sciencedirect.com/science/article/pii/S0045782516304042?dgcid=raven{_}sd{_}recommender{_}email

[55] E. Bar-Kochba, J. Toyjanova, E. Andrews, K. S. Kim, C. Franck, A Fast Iterative Digital Volume Correlation Algorithm for Large Deformations, Experimental Mechanics 55 (1) (2015) 261–274. doi:10.1007/s11340-014-9874-2.

[56] M.-M. Vaeyens, A. Jorge-Peñas, J. Barrasa-Fano, C. Steuwe, T. Heck, P. Carmeliet, M. Roeffaers, H. Van Oosterwyck, Matrix deformations around angiogenic sprouts correlate to sprout dynamics and suggest pulling activity, Angiogenesisdoi:10.1007/s10456-020-09708-y. URL http://link.springer.com/10.1007/s10456-020-09708-y

[57] D. Garcia, Robust smoothing of gridded data in one and higher dimensions with missing values, Computational Statistics & Data Analysis 54 (4) (2010) 1167–1178. doi:10.1016/J.CSDA.2009.09.020. URL https://www.sciencedirect.com/science/article/pii/S0167947309003491?via{%}3Dihub

[58] N. Otsu, A Threshold Selection Method from Gray-Level Histograms, IEEE Transactions on Systems, Man, and Cybernetics 9 (1) (1979) 62–66. doi: 10.1109/TSMC.1979.4310076. URL http://ieeexplore.ieee.org/document/4310076/

[59] Qianqian Fang, D. A. Boas, Tetrahedral mesh generation from volumetric binary and grayscale images, in: 2009 IEEE International Symposium on Biomedical Imaging: From Nano to Macro, IEEE, 2009, pp. 1142–1145. doi:10.1109/ISBI.2009.5193259. URL http://ieeexplore.ieee.org/document/5193259/

[60] D. Ambrosi, Cellular Traction as an Inverse Problem, SIAM Journal on Applied Mathematics 66 (6) (2006) 2049–2060. doi:10.1137/060657121. URL http://epubs.siam.org/doi/10.1137/060657121

[61] D. Ambrosi, A. Duperray, V. Peschetola, C. Verdier, Traction patterns of tumor cells, Journal of Mathematical Biology 58 (1-2) (2009) 163–181. doi:10.1007/s00285-008-0167-1.

[62] P. Carmeliet, R. K. Jain, Molecular mechanisms and clinical applications of angiogenesis, Nature 473 (7347) (2011) 298–307. doi:10.1038/nature10144. URL http://www.nature.com/articles/nature10144

[63] T. Heck, M. M. Vaeyens, H. Van Oosterwyck, Computational Models of Sprouting Angiogenesis and Cell Migration: Towards Multiscale Mechanochemical Models of Angiogenesis, Mathematical Modelling of Natural Phenomena 10 (1) (2015) 108–141. doi:10.1051/mmnp/201510106. URL http://www.mmnp-journal.org/10.1051/mmnp/201510106

[64] T. Lee, R. Kashyap, C. Chu, Building Skeleton Models via 3-D Medial Surface Axis Thinning Algorithms, CVGIP: Graphical Models and Image Processing 56 (6) (1994) 462–478. doi:10.1006/CGIP.1994.1042. URL https://www.sciencedirect.com/science/article/pii/S104996528471042X

[65] R. van den Boomgaard, R. van Balen, Methods for fast morphological image transforms using bitmapped binary images, CVGIP: Graphical Models and Image Processing 54 (3) (1992) 252–258. doi:10.1016/1049-9652(92)90055-3. URL https://www.sciencedirect.com/science/article/pii/1049965292900553

[66] B. Zhang, J. Zerubia, J.-C. Olivo-Marin, Gaussian approximations of fluorescence microscope point-spread function models, Applied Optics 46 (10) (2007) 1819. doi:10.1364/AO.46.001819. URL https://www.osapublishing.org/abstract.cfm?URI=ao-46-10-1819

[67] DIPimage - DIPimage & DIPlib. URL http://www.diplib.org/dipimage

[68] A. Jorge-Peñas, H. Bové, K. Sanen, M.-M. Vaeyens, C. Steuwe, M. Roeffaers, M. Ameloot, H. Van Oosterwyck, 3D full-field quantification of cell-induced large deformations in fibrillar biomaterials by combining non-rigid image registration with label-free second harmonic generation, Biomaterials 136 (2017) 86–97. doi:10.1016/j.biomaterials.2017.05.015. URL https://ac.els-cdn.com/S0142961217303277/1-s2.0-S0142961217303277-main.pdf?_tid=6d03138e-fd25-11e7-b858-00000aacb35d&acdnat=1516372464_9e13203f92977e35974bc5e7e741f62b https://linkinghub.elsevier.com/retrieve/pii/S0142961217303277

